# Analytical Choices Drive Toxicogenomic Potency Estimates: A Systematic Evaluation of Transcriptomic Points of Departure

**DOI:** 10.64898/2026.05.27.728212

**Authors:** Imke B. Bruns, Dayna R. Schultz, Emmanuel Demuynck, Friedel Dewulf, Ioannis Theologidis, Steven J. Kunnen, Lukas S. Wijaya, Ilias Frydas, Nafsika Papaioannou, Elisavet Renieri, Thanasis Papageorgiou, Dimosthenis Sarigiannis, Kyriaki Machera, Birgit Mertens, Jana Asselman, Carsten Weiss, Bob van de Water, Giulia Callegaro

**Affiliations:** Division of Drug Discovery and Safety, Leiden Academic Centre for Drug Research, Leiden University, Leiden, the Netherlands; Aristotle University of Thessaloniki, Thessaloniki, Greece; HERACLES Research Center on Health and the Exposome, Center for Interdisciplinary Research and Innovation, Aristotle University of Thessaloniki, Greece; Scientific Direction of Chemical and Physical Health Risks, Sciensano, Brussels, Belgium; Department of In Vitro Toxicology and Dermato-Cosmetology, Vrije Universiteit Brussel, Brussels, Belgium; Blue Growth Research Lab, Ghent University, Ghent, Belgium; Laboratory of Toxicological Control of Pesticides, Scientific Directorate of Pesticides’ Control & Phytopharmacy, Benaki Phytopathological Institute, Kifissia, Greece; National Hellenic Research Foundation, Athens, Greece; Institute of Biological and Chemical Systems - Biological Information Processing, Karlsruhe Institute of Technology, Karlsruhe, Germany

**Keywords:** Transcriptomic point of departure, Benchmark Concentration, Workflow Variability, In Vitro Transcriptomics, Next Generation Risk Assessment

## Abstract

Omics technologies are increasingly integrated into next-generation risk assessment, yet quantitative toxicogenomics outcomes remain highly dependent on analytical choices, motivating a systematic evaluation of how bioinformatics workflows influence hazard characterization and transcriptomic Points of Departure (tPOD). Here, we applied five independent transcriptomics pipelines to a shared dataset of RPTEC-TERT1 kidney cells exposed to cisplatin across multiple concentrations and timepoints, comparing effects of pre-processing, benchmark concentration modeling, and pathway-based interpretation strategies. Across workflows, substantial variability was observed in gene-level benchmark concentrations (BMCs), primarily driven by differences in normalization, filtering, and especially the modeling software used. Despite this variability, convergence increased at later timepoints as transcriptional responses strengthened, with 24 h consistently identified as the most sensitive timepoint at the gene level. Aggregation of gene-level BMCs into pathway-based metrics reduced variability but did not eliminate it, with pathway definition emerging as a major determinant of sensitivity estimates. Notably, distinct pathway resources showed minimal gene overlap, and smaller, biologically coherent gene sets (e.g., co-expression modules and biomarker panels) produced lower and less dispersed BMCs compared with broader pathway annotations. Furthermore, direct modeling of pathway activity scores yielded systematically different sensitivity estimates relative to median-based aggregation, with method-dependent conservativeness influenced by pathway coverage and response strength. Overall, our findings demonstrate that both analytical workflow design and pathway selection critically shape toxicogenomic-derived potency estimates, highlighting the need for harmonized, transparent methodologies to enable robust application of transcriptomics in chemical safety assessment and regulatory decision-making.

## Introduction

There is increasing evidence suggesting that integrating omics technologies, particularly transcriptomics, into chemical risk assessment (CRA) frameworks can enhance hazard characterization and support the derivation of points of departure (PODs) (Mezencev and Subramaniam 2019; del Giudice et al. 2023). Transcriptomics can provide mechanistic insight into stressor mode of action when gene expression changes are translated into functional biological knowledge, particularly within tiered testing strategies that included targeted and organotypic assays (Thomas et al. 2019; Rogers et al. 2025). Accordingly, several frameworks have been developed to guide the use of omics data in CRA and to distinguish adverse from adaptive biological responses (Johnson et al. 2022).

A key application of transcriptomics in hazard assessment is the derivation of transcriptomic PODs (tPODs) (NTP 2018). Standard tPOD workflows typically include quality control and normalization, identification of concentration-responsive genes, benchmark concentration (BMC) modeling of individual genes, post-model quality filtering, and summarization of gene-level BMCs into transcriptome wide or pathway-level tPOD (U.S. Environmental Protection Agency 2012; Crizer et al. 2021; Feshuk et al. 2023; Reardon et al. 2023; Costa et al. 2024; O’Brien et al. 2025). Gene-level BMCs can be summarized using distribution-based approaches, such as percentile methods or the lowest consistent response dose, or gene set-based approaches based on the lowest median BMC among responsive pathways.

Despite the clear advantages and progress of these approaches, several sources of variability must be addressed to strengthen the reliability and regulatory acceptance of tPODs (Aurisano et al. 2023; Gant et al. 2023). These include technical and biological variability in transcriptomic data, differences in preprocessing and modeling workflows, the aggregation of gene-level BMCs into pathway- or distribution-based tPODs, and the challenge of distinguishing adaptive from toxicologically relevant responses (Costa et al. 2024). In addition, experimental design choices, particularly exposure duration in *in vitro* studies, may strongly influence BMC and tPOD derivation, as supported by recent evidence of time-dependent variation in BMC values (Tennekes and Sánchez-Bayo 2013; Aguayo-Orozco et al. 2018; Carpi et al. 2024; Harrill, Everett, Haggard, Bundy, et al. 2024).

Several of these issues have been investigated thoroughly and, in some cases, there is a set path to potentially resolve them. Standardized reporting frameworks have been developed and tested (OECD 2023 Nov 22), although their reproducibility across multiple analysts working independently on the same dataset has yet to be demonstrated. Comprehensive examples and guidelines for recommended pipeline steps have been published (Reardon et al. 2023; Harrill, Everett, Haggard, Word, et al. 2024; O’Brien et al. 2025), however it is still unclear which specific processing steps exert the greatest influence on tPOD determination at either the gene- or pathway-level. Recent studies have evaluated distribution-based versus gene set-based tPOD approaches (Costa et al. 2024), suggesting broadly similar overall estimates. However, uncertainties remain regarding the robustness of these assessments, particularly with respect to gene set composition and variability in individual tPOD estimates (Beebe-Wang et al. 2022). Further research is needed to optimize gene set selection for tPOD derivation, including the use of annotation-based pathways (e.g. KEGG, WikiPathways, HALLMARK) (Kanehisa and Goto 2000; Liberzon et al. 2015; Slenter et al. 2018), curated transcriptomic signatures (e.g. GENOMARK, TGx-DDI) (Li et al. 2015; Ates et al. 2018), or data-driven gene co-expression modules (van Kessel et al. 2025 Nov 20).

In this study, we addressed the points above by applying five independent transcriptomic analysis workflows, representative of current practice in the field, to derive tPODs. All workflows were applied to the same transcriptomic dataset, generated in human immortalized renal proximal tubule epithelial cells RPTEC-TERT1 following exposure to the clinically relevant nephrotoxicant cisplatin across nine concentrations and six timepoints. The OECD Omics Reporting Framework (OORF) was used to systematically document technical choices, including normalization and downstream modeling steps (OECD 2023 Nov 22). Using this framework, we examined how variability introduced during data processing and modeling affects gene-level BMCs, gene set-derived median BMCs, and study-level tPODs. We show that differences in data preprocessing and filtering contribute substantially to variability prior to modeling, with the most pronounced differences arising from the use of different BMD modeling software. Post-model filtering criteria can partially harmonize gene-level results across the tested workflows. Temporal analysis revealed a time-dependent convergence in transcriptomic responses, with later timepoints showing increased coherence in derived BMCs and tPODs. We further demonstrate that pathway-level outcomes depend strongly on pathway definition: large, curated pathways showed greater heterogeneity, while system-specific WGCNA gene co-regulation modules and biomarker gene sets yielded more consistent results. Finally, comparisons of median-based and pathway score-based modeling highlighted complementary strengths and limitations, underscoring the importance of pathway-level coverage when deriving pathway-based tPODs. Together, these findings provide practical insights into methodological trade-offs and inform best practices for tPOD derivation in Next-Generation Risk Assessment (NGRA)-relevant applications.

## Materials and Methods

### Experimental design

RPTEC/TERT1 cells were maintained and differentiated in a 1:1 mixture of DMEM (Thermo Fisher Gibco, 11966-025) and Ham’s F-12 (Thermo Fisher Gibco, 21765-029), yielding a final glucose concentration of 5 mM. The medium was supplemented with 2 mM GlutaMAX (Thermo Fisher, 35050038), 10 ng/mL epidermal growth factor (Merck, E9644), 36 ng/mL hydrocortisone (Sigma-Aldrich, H0888), 5 mg/mL insulin, 5 mg/mL transferrin, 5 ng/mL selenium (Sigma-Aldrich, I1884), and 100 U/mL penicillin with 100 mg/mL streptomycin (Thermo Fisher Gibco, 15070-063). Cells were cultured at 37 °C in a humidified atmosphere with 5% CO2. For exposures, cells were seeded into 96-well plates at 25,000 cells per well and cultured for 14 days to achieve a mature proximal tubule phenotype, as described by Aschauwer *et al*. (Aschauer et al. 2015). The medium was refreshed one day prior to treatment. At t=0, cells were exposed to cisplatin in a single-exposure design at concentrations of 0.1, 0.5, 1, 2.5, 5, 10, 20, 30 and 50 µM for 4, 8, 16, 24, 48 or 72 hours. Untreated RPTEC/TERT1 cells served as controls. Cisplatin was obtained from the Leiden University Medical Center (LUMC) as a 1 mg/mL solution in 0.9% NaCl (9 mg/mL). Following exposure, cells were washed once with DPBS lacking Ca2+ and Mg2+ and lysed in 50 uL of lysis buffer (1:1 DPBS:TempO-Seq lysis buffer, BioSpyder) prepared at 2x and mixed immediately before use. Plates were sealed with aluminum sealing film (Greiner Bio-One, 676090), incubated for 15 min at room temperature, and stored at −80 °C. Prior to shipment, lysates were thawed and transferred to 96-well conical-bottom plates (Thermo Fisher, 249662), resealed with aluminum film, returned to −80 °C, and shipped on dry ice to BioClavis (UK). Samples were processed using the TempO-Seq platform with the human whole transcriptome v1.2 probe panel. In May 2023, the probe panel was realigned to the reference genome (GRCh38) using a 96% sequence identity threshold. Standard attenuators were applied according to the manufacturer’s protocol.

### General Workflow

#### Quality Control and Normalization

Each partner followed their own specific pipeline based on pre-existing, laboratory-specific protocols, and all analyses adhered to a shared conceptual framework for deriving BMCs from transcriptomics data adapted from O’Brien *et al*. (O’Brien et al. 2025) (**Figure 1A**). First, quality control (QC) was performed on the raw aligned TempO-Seq counts, including filtering of low-quality probes and exclusion of samples with low library size or poor replicate correlation. After QC, gene-level expression values were derived by aggregating probe counts and performing normalization.

**Figure 1.**
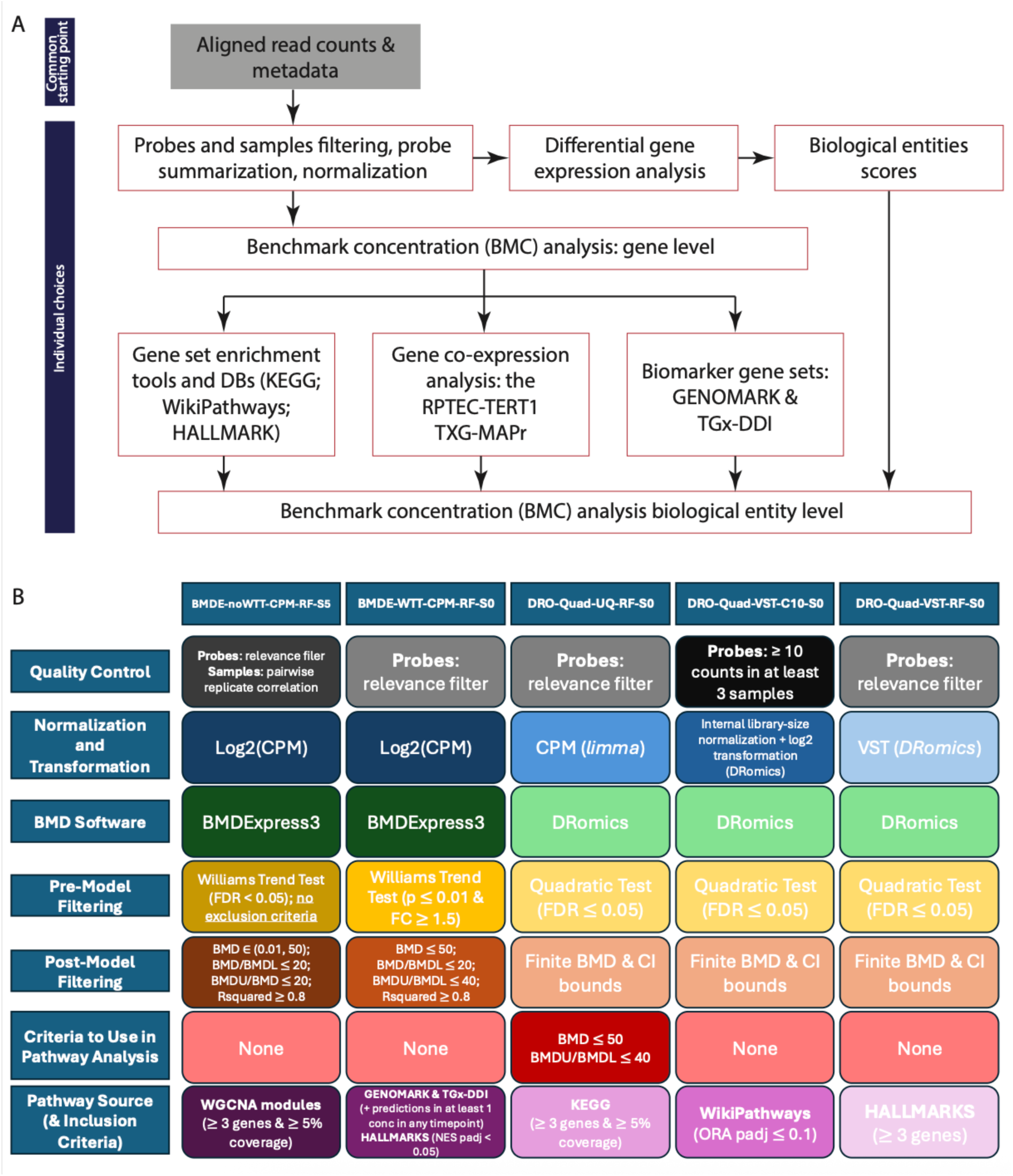
Schematic overview of the analysis workflow. All partners began from the same aligned read counts to minimize variability in preprocessing. From these counts, each partner then carried out their own quality control, normalization, benchmark concentration (BMC) modeling, and pathway interpretation (**A**). Key differences across pipelines included the quality control and normalization steps, the BMD software used, the filtering steps applied, and the pathway sources used (**B**).

#### Determine Gene Set Scores

In some cases, log2 fold changes (log2FC) were calculated using DESeq2 (Love et al. 2014). These log2FC values were also used to generate pathway-level activity scores: either by uploading the data to the RPTEC/TERT1 TXG-MAPr tool to obtain eigengene scores based on a predefined WGCNA network (van Kessel et al. 2025 Nov 20), or by performing gene set enrichment analysis (GSEA) to calculate normalized enrichment scores (NES) (Subramanian et al. 2005).

#### Benchmark Concentration Modeling

Benchmark concentration modeling was applied to normalized gene expression data, yielding gene-level BMCs (Yang et al. 2007; Larras et al. 2018). Due to inherent challenges in summarizing transcriptomic-based BMC data into a single tPOD, multiple strategies were implemented and compared. Specifically, we evaluated four gene-level distribution-based approaches for calculating tPODs: the fifth percentile, the first mode, the 25^th^ lowest ranked gene, and the lowest consistent response dose (LCRD) (Farmahin et al. 2017). In addition, gene-level BMCs were aggregated at the pathway level, and the median BMC was used to derive a pathway-based tPOD.

#### Distribution-Based tPODs

The fifth percentile method defines the tPOD as the BMC of the gene nearest to the 5^th^ percentile of the gene-level BMC distribution (Farmahin et al. 2017; Reardon et al. 2021; Reardon et al. 2023). The 25^th^ ranked gene approach orders all gene-level BMCs and selects the BMC of the 25^th^ gene (Reardon et al. 2021; Matteo et al. 2023; Reardon et al. 2023). Conversely, the first mode identifies the earliest peak in the BMC density curve using a second-derivative inflection-point criterion (Pagé-Larivière et al. 2019). The Kneedle algorithm was applied to identify the point of maximum curvature (“knee-point”) in the ordered BMC distribution (Satopää et al. 2011). Finally, the LCRD is defined as the lowest BMC within the largest group of consecutive BMCs for which the ratio between adjacent values does not exceed a factor of 1.66, forming the consistent group of BMCs (CRGB). Following the adaptation described by Reardon *et al*., the largest qualifying group is considered rather than the earlies group with subsequent increasing BMCs (Crizer et al. 2021; Reardon et al. 2023).

In addition, to quantify uncertainty in the derived tPOD estimates, the gene-level BMC distributions were bootstrapped (n = 1000 iterations). In each bootstrap iteration, BMC values were resampled with replacement while preserving the original sample size. For each resampled distribution, tPODs were recalculated, and the resulting distributions were summarized using the median and corresponding 95% confidence intervals.

#### Pathway-Based tPODs

To interpret the responses at a higher biological level, genes were grouped into predefined gene sets (e.g. KEGG (Kanehisa and Goto 2000), HALLMARK (Liberzon et al. 2015), WikiPathways (Slenter et al. 2018), WGCNA modules (van Kessel et al. 2025 Nov 20), or the TGx-DDI and GENOMARK biomarker panels (Li et al. 2015; Ates et al. 2018)), and pathway-level BMCs were calculated as the median BMC of all concentration-responsive genes in each set. In parallel, BMC modeling was also applied directly to pathway activity scores (eigengene scores or normalized enrichment scores) to enable comparisons between tPODs derived from gene-level aggregation into pathways and those obtained from direct pathway-level modeling (**Figure 1A**).

To evaluate the effect of sample size on correlations of HALLMARK pathway median BMCs between workflows, we performed a subsampling analysis of gene-level BMCs. For each workflow pair and timepoint, the number of overlapping pathways contributing to the correlations was determined. Gene-level BMCs were then randomly subsampled 2,000 times, each time selecting the same number of genes as the number of overlapping pathways used in the corresponding pathway-level correlation. Pearson correlations derived from these subsampled gene-level BMCs were subsequently compared to the observed pathway-level median BMC correlations.

#### Details on Specific Workflows

Details on specific workflows followed by each partner are described in **Supplemental Table 2** and based on selected elements of the OORF (OECD 2023 Nov 22). Key differences are also highlighted in **Figure 1B**, and full methodological descriptions are provided in the Supplementary Information. All preprocessing code and partner-specific analysis scripts are accessible in the corresponding GitHub folders and have been archived on Zenodo to provide a permanent, citable record (DOI: https://doi.org/10.5281/zenodo.20413136).

To facilitate transparent comparison of analytical approaches, each workflow was assigned a structured identifier reflecting key methodological choices. Workflow names encode the BMD software used, prefilter application, normalization method, probe-level quality control filter, and sample exclusion. For readability abbreviated identifiers are used throughout the main text, while the full workflow names are provided in **Table 1**.

**Table 1.** Overview of the analytical workflows and naming convention. Abbreviated workflow identifiers are used in the main text to enhance readability, whereas full workflow names are presented in figures and tables to ensure clarity and methodological transparency.

| Abbreviated workflow identifiers | Full workflow name | BMD software | Prefilter | Normalization | Probe QC | Number of samples removed |
| --- | --- | --- | --- | --- | --- | --- |
| BMDE-noWTT | BMDE-noWTT-CPM-RF-S5 | BMDEExpress 3 [BMDE] | None [noWTT] | Counts per million [CPM] | Relevance filter [RF] | 5 samples removed [S5] |
| BMDE-WTT | BMDE-WTT-CPM-RF-S0 | BMDEExpress 3 [BMDE] | Williams Trend Test [WTT] | Counts per million [CPM] | Relevance filter [RF] | 0 samples removed [S0] |
| DRO-UQ | DRO-Quad-UQ-RF-S0 | DRomics [DRO] | Quadratic test [Quad] | Upper quartile [UQ] | Relevance filter [RF] | 0 samples removed [S0] |
| DRO-VST-C10 | DRO-Quad-VST-C10-S0 | DRomics [DRO] | Quadratic test [Quad] | Variance stabilization transformation [VST] | Retained probes if they had $\geq 10$ counts in $\geq 3$ samples [C10] | 0 samples removed [S0] |
| DRO-VST-RF | DRO-Quad-VST-RF-S0 | DRomics [DRO] | Quadratic test [Quad] | Variance stabilization transformation [VST] | Relevance filter [RF] | 0 samples removed [S0] |

### Reporting Framework for Benchmark Concentration Analysis

To standardize and facilitate the interpretation of concentration-response modeling results across datasets and workflows, we developed a reproducible reporting framework implemented as an R Markdown document. The framework accepts multiple input types, including normalized expression values (i.e., counts per million), log2FC, and pathway-level scores (e.g. BMCs derived from WGCNA eigengene scores or normalized enrichment scores), depending on the analysis performed. For each analysis type, users specify relevant metadata and model output files, after which a series of standardized visualizations and summaries are generated.

The framework is structured into four analytical components that summarize and visualize results across different methods. The framework itself does not perform BMC modeling but instead integrates and presents pre-existing analysis outputs in a standardized and interactive format. First, an initial data overview is provided using principal component analysis (PCA) to visualize global transcriptional patterns and assess concentration-dependent separation among samples. Second, gene-level BMCs are summarized to report the number of concentration-responsive genes and interactive accumulation plots. Based on these uploaded results, distribution-based tPODs are also derived, including the 5^th^ percentile BMC, the first mode, the 25^th^ ranked BMC and the LCRD of the distribution. Third, pathway-level tPODs derived from gene aggregation are summarized using interactive tables, including reported BMC estimates and confidence intervals, with search functionality to enable targeted pathway exploration. In addition, the distributions of gene-level BMCs within the four most responsive pathways are visualized in relation to the aggregated pathway BMC and its corresponding confidence interval. Fourth, when pathway score-based BMC modeling results are provided, these outputs are summarized to report concentration-responsive pathways, accumulation plots, and pathway-based tPODs using the same distributional approaches applied at the gene level. If any of the results are not provided, the framework automatically omits that component. All summaries are organized by timepoint, with separate tabs to facilitate interactive exploration.

The reporting framework was applied independently by each partner, and the resulting HTML reports can be found in the supplemental materials. In addition, the source code and documentation for the framework is publicly accessible via GitHub (https://github.com/vdWater-Callegaro-lab/PARC-abInitio) and Zenodo (DOI: https://doi.org/10.5281/zenodo.20413136).

### Code and Data Availability

All data analyzed during this study are available in the BioStudies repository under accession number E-MTAB-11374 and are available upon request. The code used for data processing, analysis and figure generation is publicly available on GitHub at https://github.com/vdWater-Callegaro-lab/PARC-abInitio, and a versioned snapshot of the repository has been deposited on Zenodo (DOI: https://doi.org/10.5281/zenodo.20413136) to ensure long-term accessibility. Analyses performed by each partner are documented in the corresponding partner-specific folders within the GitHub repository.

## Results

### Gene-level BMC Approaches Consistently Identify 24h as the Most Sensitive Timepoint but Yield Substantial Variation in Individual tPOD Estimates

#### Data Preprocessing Before BMC Analysis

Each workflow implemented different quality-control steps to the raw count data, including filtering of low-quality probes and, in some cases, exclusion of samples. As a result, the final number of retained genes differed across sites, with 13,040, 12,272, 12,937, 11,531 and 11,531 genes preserved for DRO-UQ, DRO-VST-C10, DRO-VST-RF, BMDE-noWTT, and BMDE-WTT, respectively. BMDE-noWTT additionally excluded five samples on quality criteria: one sample (8 h, 0.5 µM) was removed due to a low library size (<500,000 reads), evaluated after filtering low-quality probes, and four samples (16 h, 0.1 µM; 48 h 0.5 µM; 72 h 10 µM; 72 h, 20 µM) due to poor replicate correlation. Other workflows retained all samples to maintain a minimum n = 3 per replicate (**Table 2**).

**Table 2.** Data dimensions after quality control (QC) for each workflow and normalizations/transformations applied to the data before benchmark concentration modeling.

|  | Genes after QC (n) | Samples after QC (n) | Normalization and transformation |
| --- | --- | --- | --- |
| **BMDE-noWTT-CPM-RF-S5** | 11,531 | 199 | Log2(CPM) |
| **BMDE-WTT-CPM-RF-S0** | 11,531 | 204 | Log2(CPM) |
| **DRO-Quad-UQ-RF-S0** | 13,040 | 204 | Upper Quartile |
| **DRO-Quad-VST-C10-S0** | 12,272 | 204 | Log2(VST) in DRomics |
| **DRO-Quad-VST-RF-S0** | 12,937 | 204 | Log2(VST) in DRomics |

#### Number of Concentration Responsive Genes (CRG) Responding to Chemical Exposure

Selecting significantly responding genes is an important step to limit the number of features entering tPOD analyses and to remove noisy or weak concentration-response patterns. This step was implemented differently across workflows. DRO-UQ, DRO-VST-C10 and DRO-VST-RF applied a quadratic trend test with False Discovery Rate-adjusted p-value (FDR < 0.01) in DRomics as a pre-model filter, thereby restricting BMC modeling to genes that showed statistically significant concentration responsiveness. BMDE-WTT and BMDE-noWTT used the Williams Trend Test (WTT; p_adj_ < 0.05) in BMDExpress3 to identify concentration-responsive genes; however, BMDE-noWTT did not use WTT as an exclusion criterion before modeling and instead modeled all genes in the dataset and subsequently applied post-model filters to remove poor or noisy BMC fits. Following this strategy, notably, across all timepoints, only a small fraction of genes (≤ 2%) failed the pre-filter but remained after post-model filtering. Although numerically limited, these genes highlight the sensitivity-specificity trade-off inherent to the application of a prefilter. Retaining such genes may introduce a small number of false positives, whereas excluding them at the prefiltering stage could result in false negatives by removing genes with modellable concentration-response behavior (**Supplemental Figure 1**).

The number of concentration-responsive genes (CRGs) identified during the pre-filter step varied substantially across the different workflows, particularly at the early timepoints (**Table 3**). More specifically, at 4 h, DRO-UQ, DRO-VST-C10, and DRO-VST-RF identified 70, 3,618, and 295 CRGs, respectively. These results were obtained after Upper Quartile (UQ) normalization for DRO-UQ and variance-stabilizing transformation (VST) for DRO-VST-C10 and DRO-VST-RF. A key difference between the two workflows applying VST normalization lies in the probe-level filtering step. DRO-VST-C10 retained probes with ≥10 reads in at least 3 samples, whereas DRO-VST-RF applied a relevance filter for probe selection. Despite applying the same quadratic trend test within DRomics, substantial differences in the number of CRGs were observed, highlighting the influence of upstream data cleaning and normalization on CRG identification.

**Table 3.** Overview of the criteria applied to define concentration-responsive genes as a pre-filtering step for benchmark concentration (BMC) modeling, together with the number of genes classified as concentration responsive and the number retained following post-model filtering. The final column reports the mean number of concentration responsive genes and the mean number of genes retained after post-model filtering per timepoint, together with the coefficient of variation.

|  |  | **BMDE-**<br>**noWTT-**<br>**CPM-RF-S5** | **BMDE-**<br>**WTT-**<br>**CPM-RF-**<br>**S0** | **DRO-**<br>**Quad-**<br>**UQ-RF-**<br>**S0** | **DRO-**<br>**Quad-**<br>**VST-**<br>**C10-S0** | **DRO-**<br>**Quad-**<br>**VST-RF-S0** | **Mean (CV)** |
| --- | --- | --- | --- | --- | --- | --- | --- |
| **Criteria to identify concentration responsive genes** |  | Williams Trend Test (p < 0.05)<br>NOTE: no exclusion criteria | Williams Trend Test (p < 0.01 & fold filter >= 1.5) | Quadratic Test (FDR < 0.01) | Quadratic Test (FDR < 0.01) | Quadratic Test (FDR < 0.01) |  |
| **Concentration Responsive Genes (n)** | 4h | 196 | 649 | 70 | 3,618 | 295 | 966 (155.2%) |
|  | 8h | 3,304 | 2,833 | 1,782 | 5,123 | 2,347 | 3078 (41.4%) |
|  | 16h | 3,532 | 2,807 | 2,183 | 3,086 | 2,866 | 2895 (16.9%) |
|  | 24h | 8,226 | 6,477 | 4,576 | 5,801 | 5,298 | 6076 (22.9%) |
|  | 48h | 7,273 | 5,593 | 4,743 | 6,049 | 5,581 | 5848 (15.8%) |
|  | 72h | 8,895 | 6,401 | 5,245 | 6,562 | 6,028 | 6626 (20.6%) |
| **Applied Post Model Filters** |  | (1) BMD ≥ 0.01 & BMD ≤ 50; (2) BMCU/BMC ≤ 20; (3) BMC/BMCL ≤ 20; (4) Rsquared ≥ 0.8 | (1) BMC < 50 uM; (2) BMCU/BMCL ≤ 40; (3) BMC/BMCL ≤ 20; (4) Rsquared ≥ 0.8 | (1) Finite BMC point estimates & CI bounds | (1) CI within range of tested doses | (1) Finite BMC point estimates & CI bounds |  |
| **Retained genes after Post Model Filters (n)** | 4h | 76 | 42 | 66 | 2,002 | 268 | 491 (173.1%) |
|  | 8h | 701 | 695 | 1,705 | 2,731 | 2,004 | 1567 (55.9%) |
|  | 16h | 1,591 | 1,552 | 2,080 | 2,423 | 2,357 | 2001 (20.6%) |
|  | 24h | 3,312 | 3,149 | 4,272 | 4,547 | 4,471 | 3950 (16.9%) |
|  | 48h | 4,049 | 3,562 | 4,186 | 4,564 | 4,483 | 4169 (9.6%) |
|  | 72h | 5,710 | 4,061 | 4,634 | 5,073 | 4,982 | 4892 (12.4%) |

In addition, the coefficient of variation (CV) was substantially lower at later timepoints compared with 4 and 8 h, indicating reduced variability in the number of CRGs across workflows at later times. After application of post-model filtering, variability in the number of retained genes was further reduced, with counts converging to a similar range across workflows. More specifically, at 48 h the number of genes retained after post-model filtering ranged from 3,562 for BMDE-WTT to 4,564 for DRO-VST-C10, corresponding to a CV of 9.6%. At 72 h, variability increased slightly, but remained substantially lower compared to early timepoints, with retained genes ranging from 4,634 for DRO-UQ to 5710 for BMDE-noWTT (CV = 12.4%) (**Figure 2A**, **Table 3**).

**Figure 2.**
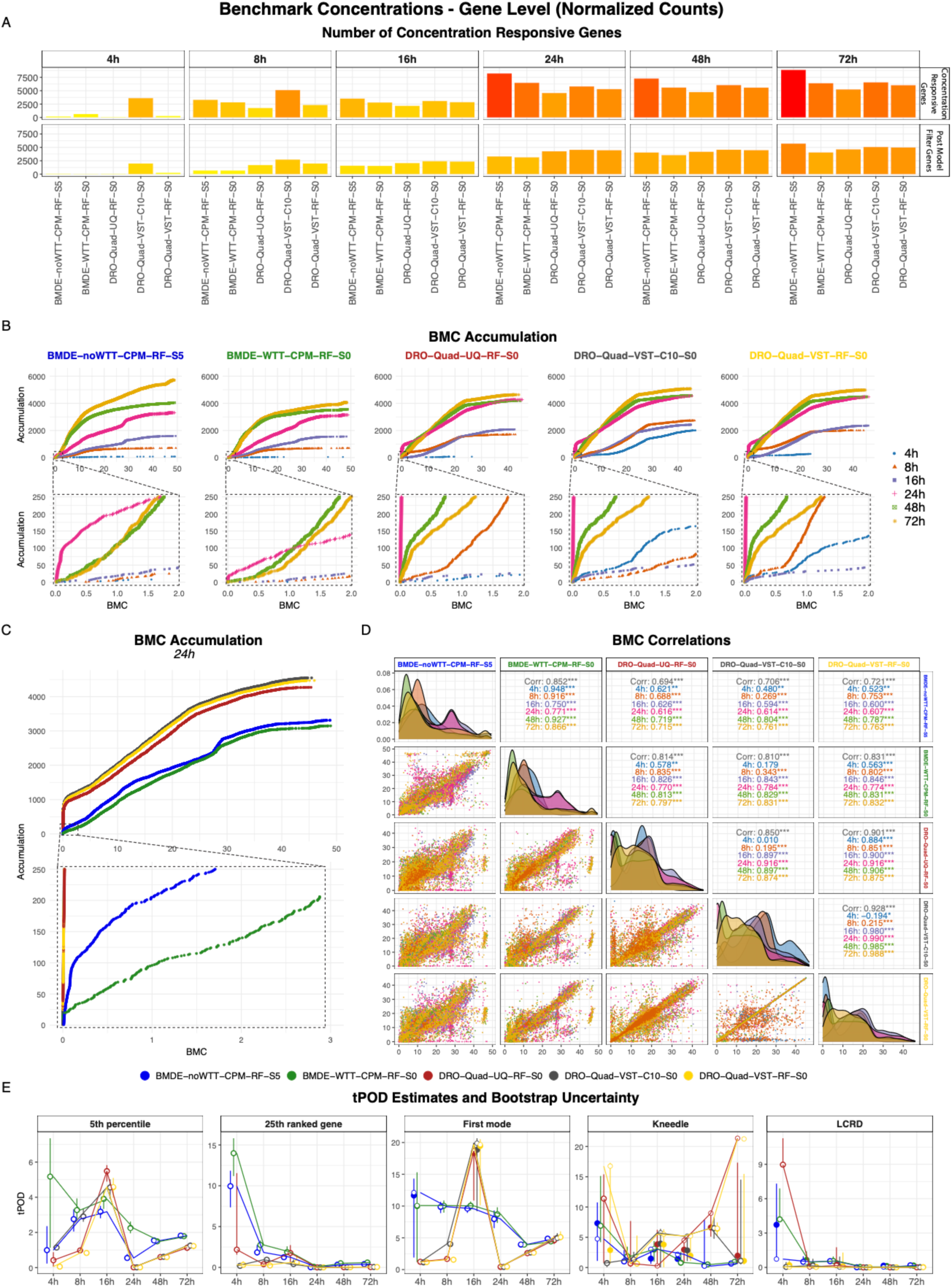
Gene-level BMC distributions and distribution-based tPODs across workflows and timepoints. Number of dose-responsive genes (CRGs) identified per workflow after application of the prefilter and after subsequent post-model filtering (**A**). Accumulation plots of all modeled genes, colored and grouped by the timepoint; inset panels show the first 250 genes with the lowest BMC values. Across all workflows, the 24 h timepoint shows the steepest initial accumulation, indicating the highest number of genes with low BMC estimates (**B**). Accumulation plots at 24 h comparing workflows, with colors indicating individual workflows (**C**). Gene-level BMC correlation plots between workflows, with colors indicating timepoints (**D**). Distribution-based tPOD estimates (5^th^ percentile, 25^th^ ranked gene, LCRD, and first mode) with uncertainty assessed by bootstrapping the BMC distributions. Filled circles represent the median bootstrapped tPOD, open circles indicate tPODs derived from the original distributions, and horizontal lines denote the bootstrapped confidence intervals (**E**).

We assessed the overlap of genes passing the pre-filter and post-model filters across workflows. The UpSet analysis of overlapping genes after both pre-filtering and post-model filtering revealed that from 16 h onward, the largest gene set consisted of genes retained across all workflows, indicating increased robustness of the transcriptional responses at later timepoints. When considering CRGs after prefiltering, BMDE-noWTT consistently exhibited the highest number of workflow-specific CRGs beyond this shared gene set. This pattern is in line with the differences in filtering criteria: DRO-UQ, DRO-VST-C10, and DRO-VST-RF applied a quadratic test with an FDR < 0.01, whereas BMDE-WTT used a Williams Trend Test (WTT) p < 0.01 combined with an absolute fold-change threshold of >= 1.5. BMDE-noWTT, in contrast, also applied WTT but used a more permissive cutoff (padj < 0.05), resulting in a larger set of genes passing the pre-filter. Conversely, although BMDE-WTT’s statistical threshold is relatively lenient, the additional fold-change requirement appears to exclude a substantial number of genes, leading to fewer genes passing the pre-filter step at later timepoints (**Supplemental Figure 2, Table 3**).

#### Gene-level BMC estimates

Gene-level BMCs were independently derived within each workflow using normalized counts and are presented as accumulation plots per timepoint (**Figure 2B**). Across all workflows, the 24 h exposure showed a steep initial increase in the accumulation curves, indicating that many genes deliver low BMC values (**Figure 2C**). This pattern is also reflected in **Table 4** and **Figure 2E**, where the 24 h timepoint consistently displayed low distribution-based tPOD estimates (5^th^ percentile, 25^th^ lowest ranked, first mode, knee-point [Kneedle], and LCRD) across workflows. At later timepoints, this early response appeared to attenuate, as evidenced by higher overall accumulation levels at 48 h and 72 h. In contrast, the earlier timepoints (4, 8, and 16 h) were characterized by more gradual increases, lower final accumulation levels, and greater variability between workflows (**Figure 2D**).

**Table 4.** Distribution-based. tPOD**s.** Summary of the distribution-based methods used to derive transcriptomic points of departure (tPODs), including the 5^th^ percentile, 25^th^ ranked gene, the first mode of the distribution, the inflection point (Kneedle) and the lowest consistent response dose (LCRD). Bootstrapped uncertainty is shown in brackets.

|  |  | BMDE-noWTT-CPM-RF-S5 | BMDE-WTT-CPM-RF-S0 | DRO-Quad-UQ-RF-S0 | DRO-Quad-VST-C10-S0 | DRO-Quad-VST-RF-S0 | Mean |
| --- | --- | --- | --- | --- | --- | --- | --- |
| 4h | 25th ranked | 10 (7.4–11.8) | 14 (11.2–15.8) | 2.2 (1.6–11.6) | 0.2 (0.1–0.5) | 0.3 (0.1–0.8) | 5.3 |
|  | 5th percentile | 1 (0.2–2.4) | 5.2 (1.1–7.4) | 0.4 (0.1–0.9) | 1.1 (1–1.3) | 0.1 (0–0.2) | 1.6 |
|  | First mode | 12.1 (1.9–14.3) | 10.1 (1.2–15.3) | 1.2 (0.9–1.5) | 1.2 (1–1.3) | 1.2 (1–1.3) | 5.2 |
|  | Kneedle | 4.7 (1.1–10.8) | 6.9 (4.2–15.1) | 11.4 (0.7–15.4) | 0.7 (0.1–1) | 16.8 (0.6–17.3) | 8.1 |
|  | LCRD | 0.7 (0.7–7.3) | 4.2 (4.2–6.9) | 9 (9–11.3) | 0 (0–0.5) | 0 (0–0.7) | 2.8 |
| 8h | 25th ranked | 1.8 (1.4–2.8) | 2.8 (2–3.3) | 0.4 (0.2–0.7) | 0.9 (0.8–1.2) | 0.2 (0.1–0.4) | 1.2 |
|  | 5th percentile | 2.8 (2.1–3.3) | 3.3 (2.8–3.9) | 1 (0.9–1.1) | 2.9 (2.6–3.3) | 0.8 (0.8–0.9) | 2.1 |
|  | First mode | 9.9 (8.9–10.7) | 10.1 (9–10.9) | 1.7 (1.6–1.8) | 4.1 (3.5–4.5) | 1.6 (1.5–1.7) | 5.4 |
|  | Kneedle | 1 (0.9–2.9) | 1.9 (0.9–3.4) | 0.7 (0.1–7.7) | 1.4 (0.9–2.6) | 0.6 (0.2–5.3) | 1.1 |
|  | LCRD | 0.5 (0.5–1) | 0.5 (0.5–1.4) | 0 (0–0.2) | 0.1 (0.1–0.8) | 0 (0–0.2) | 0.2 |
| 16h | 25th ranked | 1.2 (0.9–1.5) | 1.9 (1.5–2.6) | 1.8 (0.5–2.7) | 0.4 (0.2–1) | 0.6 (0.3–1.5) | 1.2 |
|  | 5th percentile | 3.2 (2.8–3.5) | 3.9 (3.3–4.4) | 5.5 (4.9–5.8) | 4.6 (3.9–5) | 4.6 (4–5.1) | 4.3 |
|  | First mode | 9.6 (8.9–10.2) | 10 (9.3–10.7) | 18.6 (10.9–19.3) | 19.9 (9.2–20.7) | 19.7 (10.1–20.5) | 15.6 |
|  | Kneedle | 3 (0.6–3.4) | 3.1 (1–4.4) | 0.3 (0.1–6.2) | 5.7 (0–6.4) | 5.5 (0–6.3) | 3.5 |
|  | LCRD | 0.5 (0.5–0.6) | 0.6 (0.6–1) | 0.2 (0.2–1.8) | 0 (0–0.9) | 0 (0–1.3) | 0.3 |
| 24h | 25th ranked | 0 (0–0) | 0.1 (0–0.2) | 0 (0–0) | 0 (0–0) | 0 (0–0) | 0.0 |
|  | 5th percentile | 0.5 (0.4–0.8) | 2.2 (2–2.6) | 0 (0–0) | 0 (0–0) | 0 (0–0) | 0.6 |
|  | First mode | 8 (6.5–9.3) | 8.7 (7.3–9.8) | 0.5 (0.3–0.5) | 0.4 (0.4–0.5) | 0.5 (0.4–0.5) | 3.6 |
|  | Kneedle | 2.2 (1–2.3) | 0.8 (0.3–2.6) | 4.5 (0.9–5.1) | 5.2 (1.1–5.2) | 5.2 (1.1–5.3) | 3.6 |
|  | LCRD | 0 (0–0) | 0 (0–0.2) | 0 (0–0) | 0 (0–0) | 0 (0–0) | 0.0 |
| 48h | 25th ranked | 0.3 (0.2–0.4) | 0.5 (0.4–0.6) | 0 (0–0) | 0 (0–0) | 0 (0–0) | 0.2 |
|  | 5th percentile | 1.5 (1.4–1.7) | 1.5 (1.4–1.6) | 0.6 (0.5–0.7) | 0.6 (0.5–0.8) | 0.6 (0.5–0.7) | 1.0 |
|  | First mode | 4 (3.7–4.2) | 3.9 (3.7–4.2) | 2.6 (2.4–2.8) | 2.7 (2.5–2.9) | 2.6 (2.5–2.9) | 3.2 |
|  | Kneedle | 0.2 (0.2–7.9) | 0.5 (0.4–0.7) | 9 (5.1–9.1) | 6.5 (4.7–8.3) | 6.5 (5.2–7.5) | 4.5 |
|  | LCRD | 0 (0–0.2) | 0 (0–0.3) | 0 (0–0) | 0 (0–0) | 0 (0–0) | 0.0 |
| 72h | 25th ranked | 0.4 (0.3–0.5) | 0.6 (0.5–0.8) | 0 (0–0.1) | 0.1 (0–0.1) | 0 (0–0.1) | 0.2 |
|  | 5th percentile | 1.8 (1.7–1.9) | 1.8 (1.7–1.9) | 1.1 (0.9–1.3) | 1.3 (1.1–1.4) | 1.2 (1–1.4) | 1.4 |
|  | First mode | 4.9 (4.7–5.1) | 4.4 (4.2–4.7) | 4.7 (4–5.3) | 4.9 (4.5–5.7) | 5.1 (4.6–6) | 4.8 |
|  | Kneedle | 0.7 (0.4–0.9) | 0.9 (0.5–1) | 21.4 (0.6–17.3) | 1.1 (0.5–14.5) | 21.2 (0.5–15.5) | 9.1 |
|  | LCRD | 0.2 (0.2–0.2) | 0.1 (0.1–0.3) | 0 (0–0) | 0 (0–0) | 0 (0–0) | 0.1 |

Interestingly, the shape of the accumulation plots appears to align with the modeling software used (**Figure 2B**). Workflows applying DRomics (DRO-UQ, DRO-VST-C10, and DRO-VST-RF) displayed similar curve shapes across all timepoints. Similarly, BMDExpress3-based workflows (BMDE-noWTT and BMDE-WTT) showed comparable profiles to each other, with a notable exception that BMDE-noWTT retains significantly more genes after post-model filters at 72 h, which is also shown in **Table 3**.

Despite differences in pre-processing and modeling approaches, all workflows consistently identified 24 h as the timepoint with the steepest increase in the accumulation curves at lower BMC values, indicating that many genes respond at low concentrations. When focusing on 24 h (**Figure 2C**), the primary distinction again appears to be driven by the modeling software used (DRomics versus BMDExpress3). However, when focusing on the 250 CRGs with the lowest BMCs (< 3 µM), BMDE-noWTT identified substantially more genes in the lower BMC range than BMDE-WTT, potentially reflecting the stricter prefiltering applied in the BMDE-WTT workflow. More generally, BMCs derived using DRomics tend to be lower, as reflected by the pronounced accumulation near 0 µM.

The BMC correlation matrix (**Figure 2D**) further illustrates the degree of similarity between gene-level BMC estimates across workflows. The strongest overall agreement was observed between DRO-VST-C10 and DRO-VST-RF (r = 0.928), with correlations exceeding 0.98 from the 16 h timepoint onward. A notable exception occurred at 4 and 8 h, where DRO-VST-RF consistently estimated low BMC values, while DRO-VST-C10 showed greater variability across the concentration range. Correlation between DRO-UQ and DRO-VST-RF was slightly lower (r = 0.901) yet remained high (r > 0.85) across all timepoints. In contrast, BMDE-noWTT and BMDE-WTT showed a somewhat lower overall correlation (r = 0.852), although their agreement was stronger at earlier timepoints compared with the DRomics workflows, particularly at 4 h (r = 0.948) and 8 h (r = 0.916).

Inspection of the correlation plots of gene-level BMCs generated using BMDExpress3 (BMDE-noWTT and BMDE-WTT) revealed a pronounced accumulation of BMC estimates at approximately 28 µM and 46 µM, visible as horizontal and vertical bands in the figure for 24 and 72 h, respectively. This pattern may reflect a discretization effect in the BMC estimates in BMDExpress3 rather than true biological clustering. Notably, these accumulations are not driven by the same set of genes for both BMDExpress3 workflows, as similar banding patterns are also observed in the correlation plot comparing BMDE-noWTT and BMDE-WTT. Given the experimental design, in which the tested concentration range included a relatively large interval between 30 µM and 50 µM, this pattern is likely influenced by the spacing of the tested concentrations. Examination of the fitted concentration-response models for genes contributing to the observed banding behavior revealed that the majority of genes with BMCs around 28 µM were described by an exponential 5 (Exp5) model, as shown by the vertical bands in **Supplemental Figure 3**. In addition, genes contributing to the banding observed around 46 µM were predominantly fitted by either an exponential 3 (Exp3) or a power model. These patterns are consistent between BMDE-noWTT and BMDE-WTT, suggesting that the observed banding may be associated with model fitting behavior within BMDExpress3 for these specific models (**Supplemental Figure 3**). Conversely, DRomics implements a continuous model-based approach, which avoids the derivation of discrete variables and generation of this specific artefact.

Overall, correlations were lowest at early timepoints (4-24 h), but increased markedly at longer exposures, consistently exceeding an r of 0.7 at 48 and 72 h. These findings suggest that BMC estimates at earlier timepoints are more sensitive to modeling choices, likely because transcriptional responses are still limited in intensity and less distinct due to noise. At later timepoints, when clearer and more consistent responses emerge, the results become more stable and less dependent on the specific analytical approach used.

#### Distribution-based tPODs

Gene-level tPOD estimates were obtained for each timepoint within each workflow following the BMC reporting framework (see Methods) and applying distribution-based approaches including 5^th^ percentile, 25^th^ ranked gene, first mode, knee-point (Kneedle) and lowest consistent responsive dose (LCRD) of the BMC distribution (see Methods). The LCRD consistently yielded the lowest tPOD values, except for DRO-UQ at 4h. However, LCRD values frequently fell below the lowest tested concentration of 0.1 µM, indicating that it may be overly conservative for a tPOD derivation (**Table 4**, **Supplemental Figure 4**).

To assess the uncertainty in the tPOD estimates, the BMC distributions were bootstrapped using 1,000 iterations. This analysis showed that uncertainty in tPOD derivation decreased at later timepoints, as reflected by the narrower bootstrapped confidence intervals (**Figure 2E**, **Table 4**). Among the distribution-based metrics, the Kneedle approach exhibited the greatest variability. Notably, the first mode showed a pronounced increase in variability at 16 h in workflows applying DRomics, a pattern that was consistently observed across all DRomics-based workflows. In contrast, the 25^th^ ranked gene and the LCRD produced highly consistent tPOD estimates at later timepoints, both in terms of their bootstrapped confidence intervals and their alignment across workflows.

Taken together, these findings indicate that individual gene-level BMC estimates are strongly influenced by upstream normalization choices, and most prominently, by the selected BMD modeling software. This effect was most pronounced at early timepoints, when transcriptomic responses were weak or heterogeneous, resulting in substantial variability between workflows. However, when gene-level responses were summarized into distribution-based tPODs, inter-workflow variability was markedly reduced, which was particularly evident from 24h onwards. Additionally, uncertainty associated with tPOD estimation decreased at later timepoints, as reflected by narrower bootstrapped confidence intervals, indicating increased stability.

### Reduced Correlation of Aggregated BMCs at the HALLMARK Pathway Level is Largely Driven by Feature Count

In addition to distribution-based tPODs, pathway-level aggregation was used to derive tPODs by summarizing gene-level BMCs into median pathway BMC values, allowing assessment of pathway-specific responses and potential insights into the biological mode of action. Applying a common set of HALLMARK pathways enabled direct comparison of pathway-level responses across workflows, thereby providing insight into whether aggregating functionally related genes reduced inter-workflow variability trends observed at the gene-level (Liberzon et al. 2015).

The resulting pathway-level accumulation curves (**Figure 3A**) indicated that 48 h was the most sensitive timepoint across workflows, except for DRO-VST-RF, highlighting a divergence from the gene-level analysis. Compared with the gene-level curves, the pathway-level accumulation profiles are less strongly associated with the BMD modeling software used. Interestingly, DRO-VST-RF shows a steep increase in the accumulation curve at 4 h, suggesting that some pathways responded more intensely at this early timepoint. However, this pattern is likely influenced by the absence of a pre-filter requiring a minimum percentage of pathway coverage or significant over-representation to define biologically active pathways and may be resulting from an overall minimum overlap of three genes. Specifically, at 4 h in the DRO-VST-RF workflow, only 3 of the 34 active pathways showed ≥5% gene coverage: p53 (12 genes), MTORC1 (11 genes), and TNFα signaling (15 genes). Additionally, the overrepresentation analysis identified only p53 and TNFα signaling as significant (FDR = 0.024 & FDR = 0.00357, respectively). In contrast, both DRO-UQ and BMDE-noWTT applied an additional requirement of at least 5% pathway coverage and, accordingly, did not identify any active pathways at 4 h.

**Figure 3.**
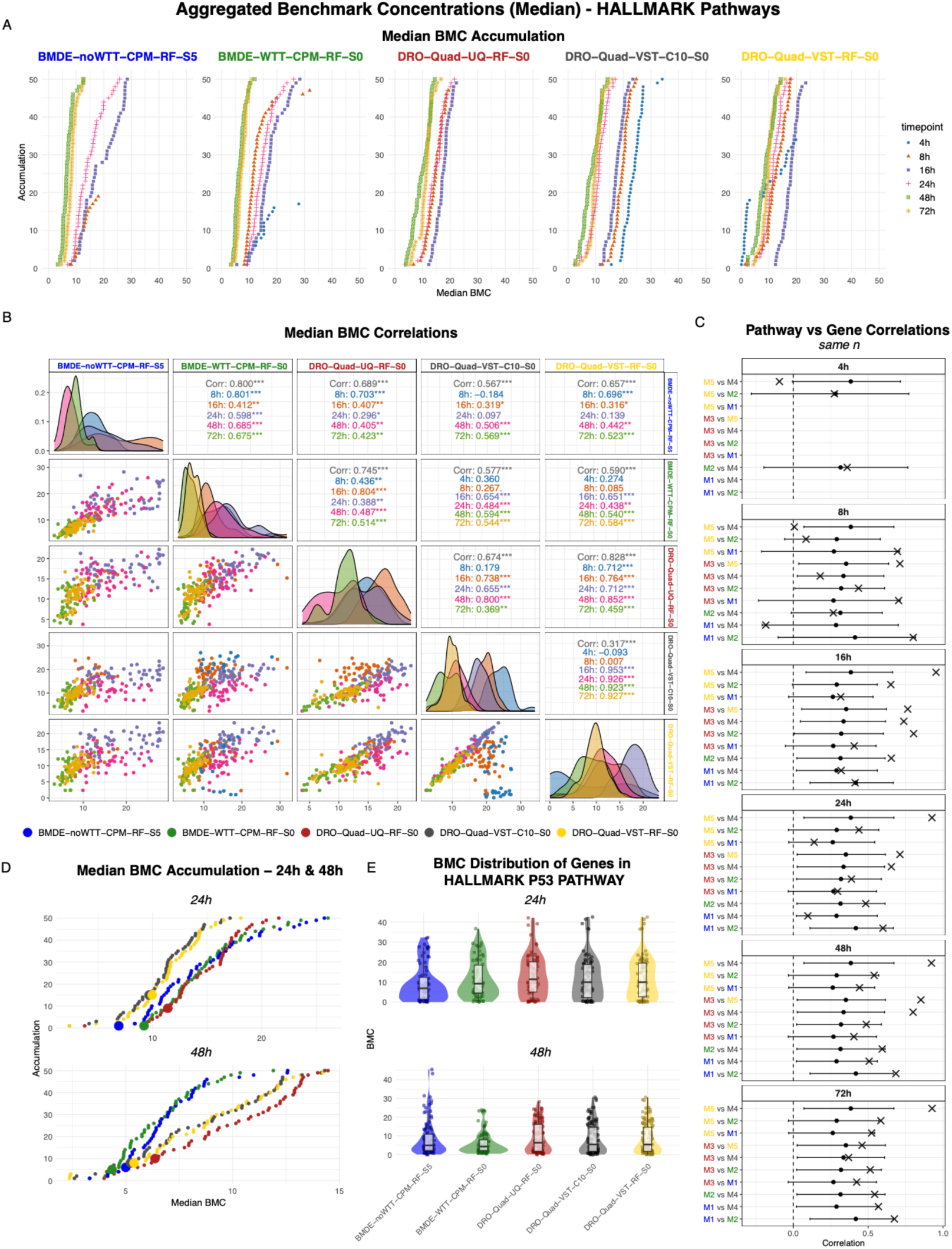
Pathway-level median BMCs derived from HALLMARK gene sets across parters and timepoints. Accumulation plots of aggregated median BMCs for HALLMARK pathways, with colors indicating timepoints (**A**). Correlation plots of pathway-level median BMCs derived from HALLMARK gene sets between workflows, with colors indicating timepoints (**B**). Observed pathway correlations versus gene-bootstrap expectations. The x marks the observed pathway correlation between workflows. Points and error bars indicate the mean and 95% confidence interval obtained by sampling the same number of genes as overlapping pathways for each workflow and timepoint. M1 = BMDE-noWTT-CPM-RF-S4, M2 = BMDE-WTT-CPM-RF-S0, M3 = DRO-Quad-UQ-RF-S0, M4 = DRO-Quad-VST-C10-S0, M5 = DRO-Quad-VST-RF-S0, (**C**). Accumulation plots of HALLMARK pathways at 24 h and 48 h, shown per workflow (colors), highlighting the HALLMARK p53 pathway (**D**). Distributions of gene-level BMCs within the HALLMARK p53 pathway for each workflow at 24 h and 48 h (**E**).

Overall, the strongest agreement across workflows in pathway-level median BMCs was observed at 48 h and 72 h. While pathway-level correlations were generally lower than gene-level correlations, this reduction is partly driven by the smaller number of pathway-level data points. Notably, sampling an equivalent number of genes resulted in poorer correlations (**Figure 3C**), indicating that pathway-level aggregation provides comparatively more robust cross-workflow concordance at matched dimensionality. Furthermore, increased variability was observed among workflows using DRomics, particularly at early timepoints. This was most evident between DRO-VST-C10 and DRO-VST-RF at 4 h (r = −0.149) and 8 h (r = −0.036), whereas correlations improved substantially at later timepoints (r ≥ 0.77 from 16 h onwards). The highest overall correlation was observed between DRO-UQ and DRO-VST-RF (overall r = 0.828). Their correlations exceeded 0.7 at all timepoints except 72 h, contrasting with the gene-level analysis, where correlations consistently increased at later timepoints.

BMDE-noWTT and BMDE-WTT also showed high overall agreement (overall r = 0.800), despite lower gene-level concordance compared with DRomics-based workflows. Nevertheless, variability between BMDE-noWTT and BMDE-WTT was most pronounced at 16 h (r = 0.412) and 24 h (r = 0.598), mirroring the timepoints at which reduced correlations were observed in the gene-level analysis.

From a pathway-oriented toxicological perspective, **Figure 3D** and **E** illustrate that workflow-specific analytical choices substantially affected sensitivity estimates, even within the well-established cisplatin-responsive HALLMARK p53 pathway (**Table 5**) (Rabik and Dolan 2006; Basu and Krishnamurthy 2010; Thompson and Joy 2024). At 24 h, both BMDE-noWTT and BMDE-WTT identified the p53 pathway as the most sensitive pathway based on median BMC values, whereas workflows using DRomics identified other pathways, including but not limited to E2F targets, G2M checkpoint, Hypoxia, and TNFα signaling as more sensitive (**Supplemental Table 2**).

**Table 5.**
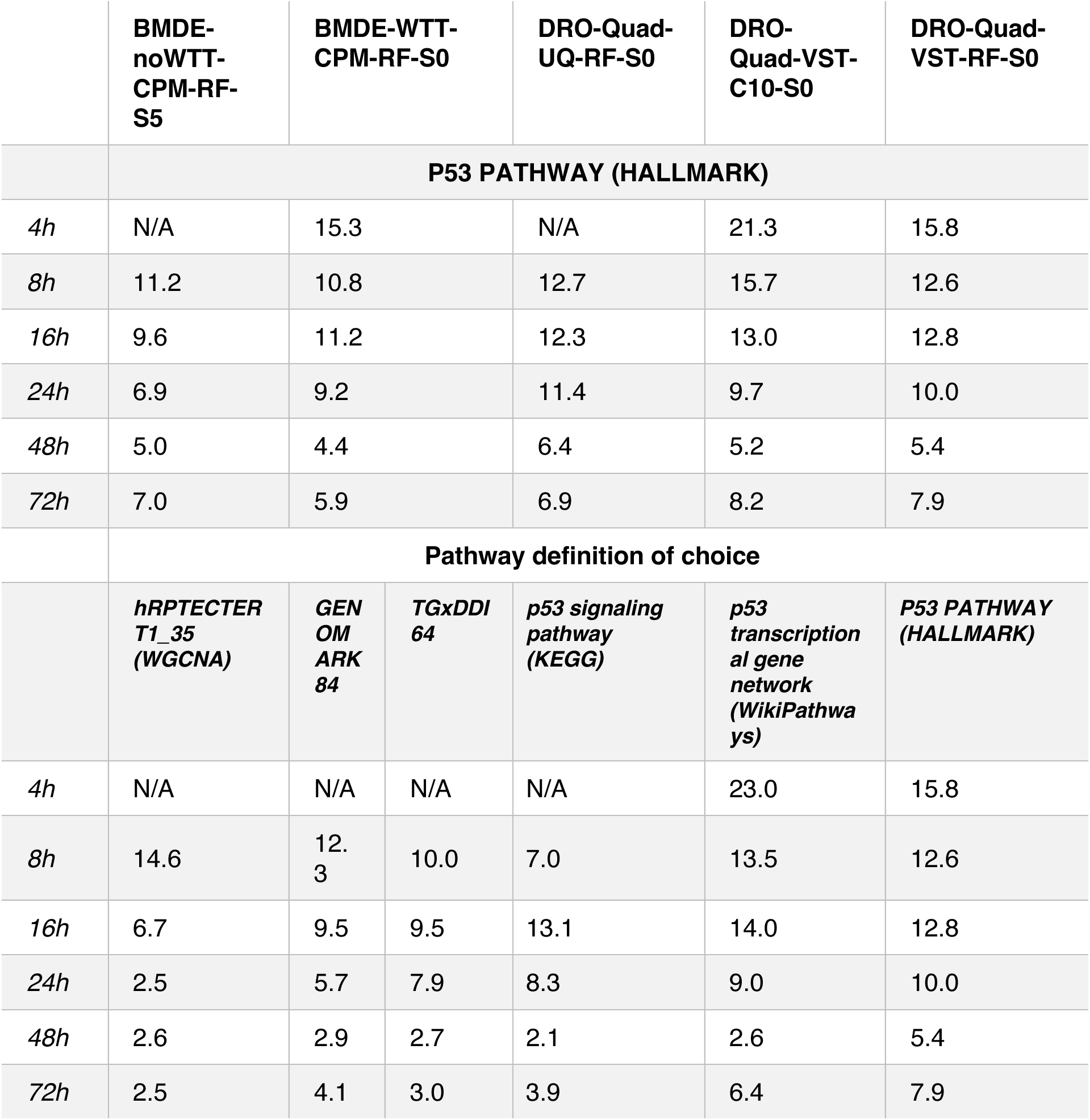
Aggregated median BMCs for p53-related pathways across workflows. The upper section presents median BMC values for the HALLMARK p53 pathway across all workflows and timepoints. The lower section summarizes median BMCs for workflow-specific p53 pathway definitions, including the KEGG p53 signaling pathway, the WikiPathways p53 transcriptional gene network, the HALLMARK p53 pathway, the hRPTECTERT1_35 WGCNA module, and the GENOMARK and TGx-DDI biomarker gene sets, as applied by DRO-UQ, DRO-VST-C10, DRO-VST-RF, BMDE-noWTT, and BMDE-WTT, respectively.

Despite these differences in ranking, aggregated median BMC values for the HALLMARK p53 pathway were comparable across workflows, ranging from 6.9 for BMDE-noWTT to 11.4 for DRO-UQ at 24 h, and converged further at 48 h (4.4 for BMDE-WTT to 6.4 for DRO-UQ). Examination of the underlying gene-level BMC distributions within the HALLMARK p53 pathway (**Figure 3D**) revealed that at 24 h, BMDE-noWTT and BMDE-WTT exhibited slightly reduced dispersion of gene-level BMCs, contributing to lower median pathway BMCs and indicating increased sensitivity. At 48 h, a stronger accumulation of genes at lower BMC values was observed across all workflows, resulting in generally lower and more comparable median BMC estimates.

Together, these findings show that although pathway-level aggregation may yield more stable cross-workflow agreement than gene-level comparisons at equivalent sample size, it does not fully remove the variability present at the gene level. Rather, a substantial proportion of this variability persists when gene-level BMCs are aggregated into median pathway BMC values.

### Variable Gene Content Across p53 Pathway Definitions Leads to Divergent tPOD Estimates, With Smaller Pathways Showing Reduced Gene-Level Dispersion

To more accurately reflect a real-world application scenario, we conducted a follow-up comparison in which each workflow selected a representative, intentionally diverse biological pathway resource for analysis. DRO-UQ used KEGG pathways (Kanehisa and Goto 2000), DRO-VST-RF selected MSigDB HALLMARK gene sets (Liberzon et al. 2015), DRO-VST-C10 utilized WikiPathways (Slenter et al. 2018), BMDE-noWTT applied their in-house RPTEC/TERT1 WGCNA modules (van Kessel et al. 2025 Nov 20), and BMDE-WTT employed the GENOMARK and TGx-DDI biomarker panels (Li et al. 2015; Ates et al. 2018), which have been previously identified as transcriptomic biomarkers of genotoxicity. This allowed for an assessment of how diverse database architectures may influence tPOD outcomes.

Given that cisplatin primarily exerts its effects through p53 signaling (Rabik and Dolan 2006; Basu and Krishnamurthy 2010; Thompson and Joy 2024), we focused specifically on p53-related signaling networks within each resource. To assess functional differences arising from the use of distinct pathway definitions and databases, we examined the overlap in gene content among the p53 pathways selected by each workflow. Specifically, the KEGG p53 signaling pathway (hsa04115; n=52 genes), the WikiPathways p53 transcriptional gene network (n=96 genes), and the HALLMARK p53 pathway (n=200 genes) were included. In addition, the RPTEC/TERT1 WGCNA module hRPTECTERT1_35 (n=33 genes), annotated as a p53 response module, was evaluated alongside the GENOMARK (n=84 genes) and TGx-DDI (n=64 genes) gene sets. Strikingly, not a single gene was shared across all pathway sets. Furthermore, only one gene (DDB2) was present in five of the selected gene sets, while most genes were unique to a single pathway database (**Figure 4A**). These findings highlight the substantial variability induced by selecting different biological entities, independent of pre-processing strategies.

**Figure 4.**
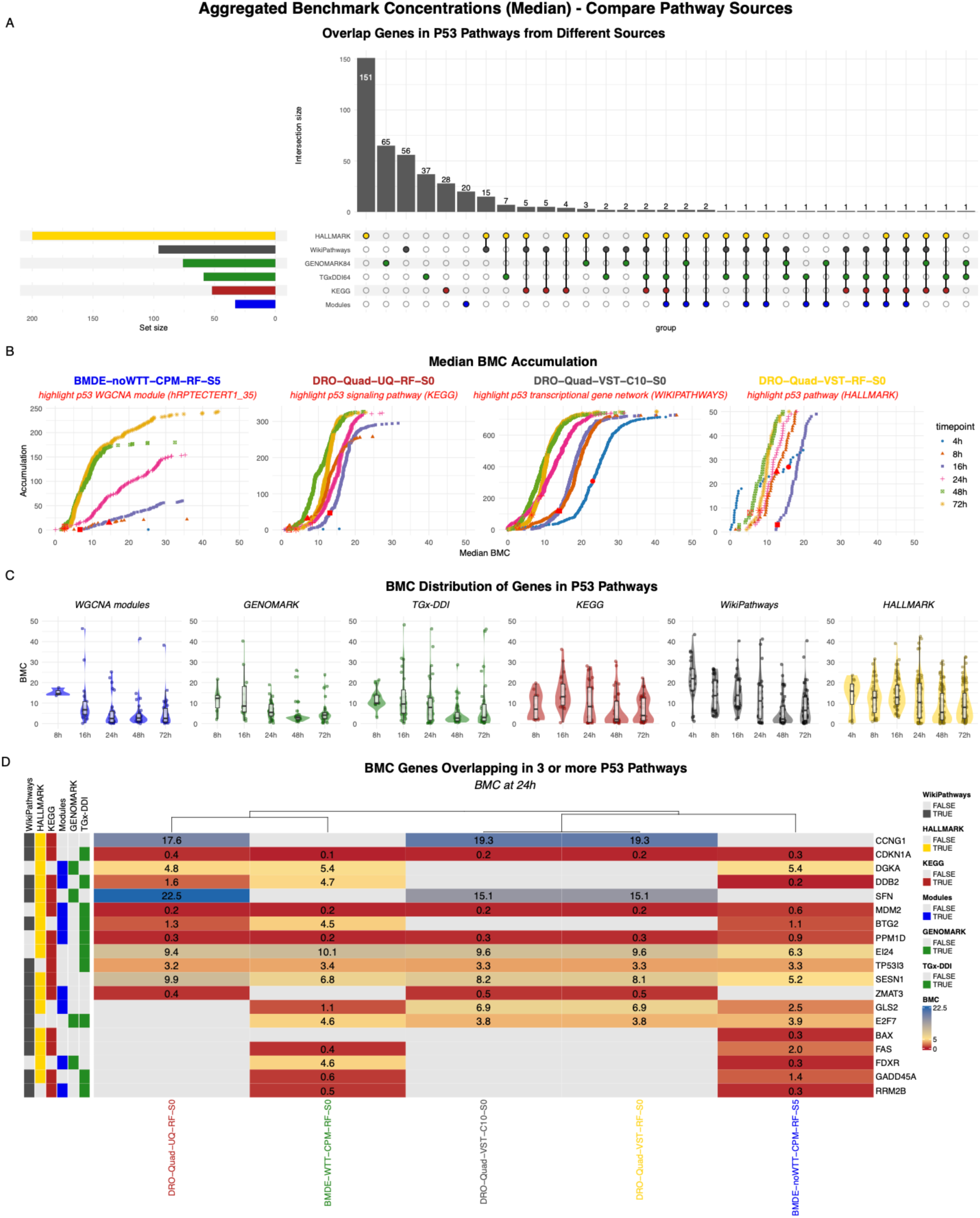
Comparison of aggregated benchmark concentrations (BMCs) derived from different p53 pathway definitions. UpSet plot showing gene overlap among p53-related pathway definitions, including the HALLMARK p53 pathway (yellow), WikiPathways p53 signaling pathway (gray), genotoxicity biomarker gene sets GENOMARK and TGx-DDI (green), the KEGG p53 signaling pathway (red), and the hRPTECTERT1_35 WGCNA module annotated as a p53 response (blue) (**A**). Accumulation plots of median BMCs aggregated for each pathway source, with p53-related pathways highlighted in red and colors indicating timepoints (B). Distributions of gene-level BMCs within each p53 pathway source (C). Heatmap of genes overlapping in three or more p53 pathway definitions. Colors indicate BMC values (red = low BMC, blue = high BMC), and annotations on the left denote the pathway sources in which each gene is included.

To assess how the choice of pathway source influences BMC determination, we compared the median BMCs derived from these different biological entities. The median BMC for the p53-related pathway was calculated for each gene set and highlighted in the accumulation plot (**Figure 4B**). Additionally, we examined the distribution of individual gene-level BMCs within each pathway and observed that larger pathways which are typically found in resources like KEGG, WikiPathways, and HALLMARK, tended to yield higher median BMCs and exhibited greater dispersion in gene-level BMC estimates (**Figure 4C**, **Table 5**). In contrast, more compact and functionally cohesive sets such as WGCNA modules and biomarker gene sets showed narrower BMC distributions and generally lower median BMC values. Notably, this was observed even in the absence of strict prefiltering. As shown previously, many genes that fail the prefilter are nonetheless retained after post-model filtering (**Supplemental Figure 1**), and the consistently low BMC values within this pathway suggest that these genes likely represent biologically relevant responses rather than noise.

Lastly, we evaluated whether genes that are more consistently associated with the p53 network exhibit lower BMC values by examining genes shared by at least three out of six p53-related pathway definitions. This analysis revealed that genes with consistently low BMCs across all workflows (PPM1D, MDM2 and CDKN1A; BMC < 1 µM for all) were identified as p53-related by at least one of the more specific pathway definitions, such as the RPTEC-TERT1 WGCNA modules or the biomarker gene sets. In contrast, genes with higher BMC values (SFN and CCNG1; > 15 µM) were predominantly annotated as p53-related in the broader pathway resources (HALLMARK, KEGG, and WikiPathways). Generally, most of the core p53 genes show comparable BMC values across workflows. For example, BMC estimates for TP53I3 ranged from 3.2 to 3.4 µM, and CDKN1A showed BMCs between 0.1 and 0.4 µM across all workflows (**Figure 4D**). Similar levels of agreement were observed for most of the remaining core p53 genes, indicating a high degree of consistency in gene-level BMCs within this core gene set.

In summary, these findings show that pathway definition influences gene set-based tPOD estimates. Larger, broadly defined pathways (e.g. KEGG, HALLMARK, and WikiPathways) exhibited greater variability in gene-level BMCs, often resulting in higher median BMC values compared to more compact system-specific (WGCNA) or curated biomarker (GENOMARK or TGx-DDI) sets. In contrast, core p53 genes shared across multiple pathway definitions showed consistent BMC values across workflows and pathways, highlighting the importance of careful pathway curation when deriving pathway-based tPODs.

### Direct Modeling of Pathway Activity Scores Alters Sensitivity Estimates Relative to Median Gene-Level BMCs

Building on the pathway-level BMC analysis, we next applied BMC modeling directly to pathway activity scores. This score-based approach was used to capture coordinated pathway responses rather than the concentration-responsive behavior of individual genes within pathways. Two complementary types of scores were evaluated: normalized enrichment scores (NES) obtained from fast GSEA of HALLMARK pathways, and eigengene scores derived from projecting log2FC values onto the existing RPTEC-TERT1 TXG-MAPr network. This approach allowed us to evaluate whether modeling pathway-level responses produces more consistent or biologically meaningful tPOD estimates than those obtained by summarizing gene-level BMCs as median values within pathways (median-based).

Figure 5A illustrates the accumulation of the pathway-level responses, with the p53 signaling pathway highlighted as an example. Based on eigengene scores (EGs) and the BMDE-noWTT workflow, the p53 pathway was not concentration responsive at 4 and 8 h but became responsive from 16 h onward. Interestingly, this pattern aligns with the median-based BMC values obtained for pathways derived from the RPTEC-TERT1 TXG-MAPr (as described previously). In the median-based analysis, the 8 h timepoint also met the inclusion threshold of at least three genes passing all model filters and a minimum of 5% pathway coverage. However, the p53 pathway at 8 h contained exactly three qualifying genes, suggesting that this signal may be an artifact of lenient filtering.

**Figure 5.**
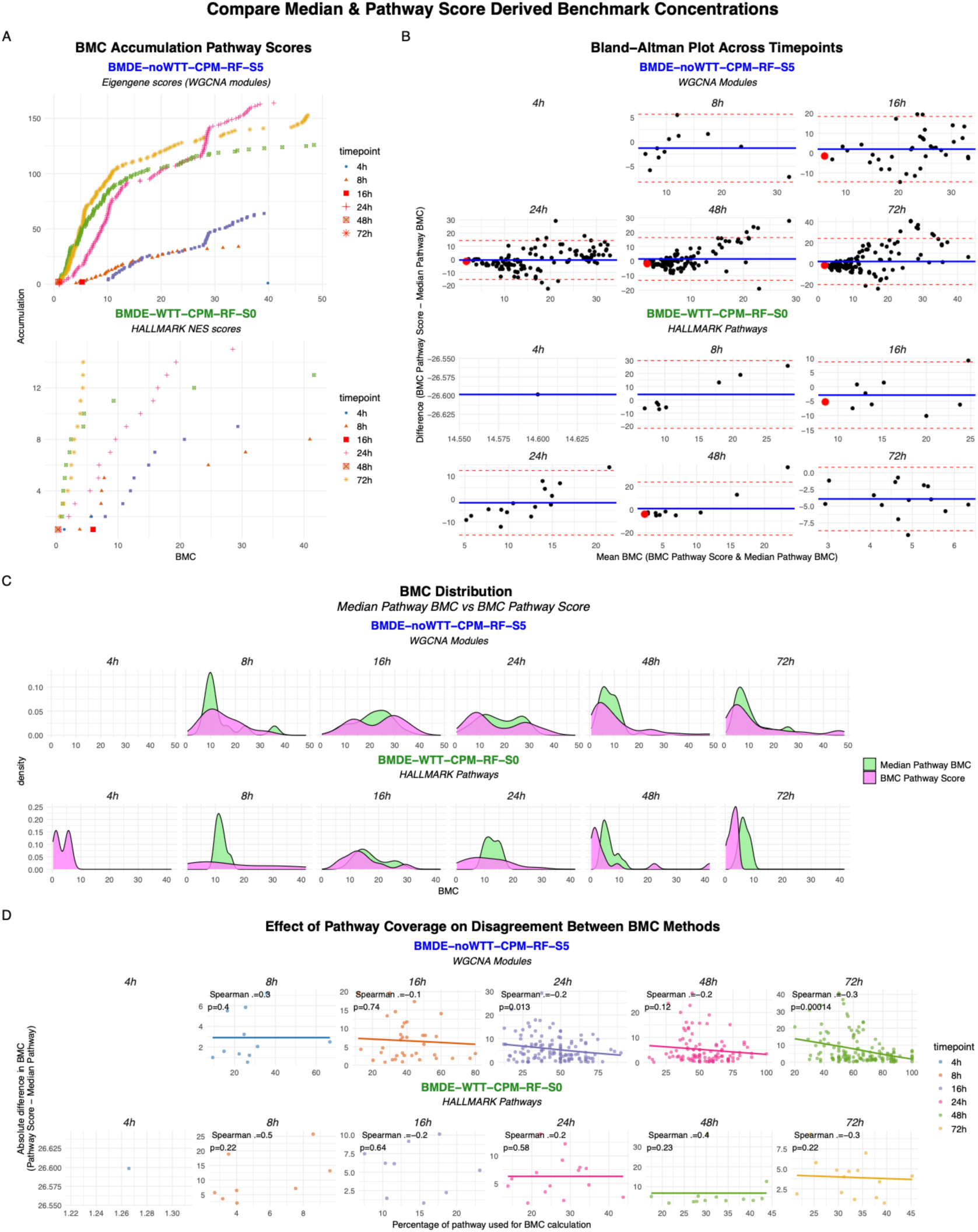
Benchmark dose modeling based on pathway scores compared with pathway-level aggregation. Accumulation plots of benchmark concentrations (BMCs) derived from direct modeling of pathway scores. BMDE-noWTT modelled WGCNA eigengene (EG) scores, while BMDE-WTT modeled normalized enrichment scores (NES) from HALLMARK pathways. Colors indicate timepoints, and the p53-related pathways (hRPTECTERT1_35 or HALLMARK p53) are highlighted in red (**A**). Bland-Altman plot comparing BMCs derived from direct modeling of pathway scores with aggregated median BMCs for the corresponding pathways or modules. The blue line indicates systematic bias between methods, and the red dotted lines represent the limits of agreement. The p53-relatede pathway or module is highlighted in red (**B**). Distributions of BMCs obtained from pathway score modeling (pink) and aggregated median pathway BMCs (green) (**C**). Relationship between pathway coverage and disagreement between BMC derivation methods. The y-axis shows the absolute difference in BMC values (pathway score-based minus aggregated median), and the x-axis shows the percentage of pathway genes contributing to the aggregated median BMC (**D**).

To compare pathway definitions based on co-expression modules with those derived from curated gene sets, we next evaluated normalized enrichment scores (NES) for the HALLMARK pathways. The BMDE-WTT workflow significantly modeled the HALLMARK p53 pathway at 16 and 48 h, but not at other timepoints. However, the score-based BMC values for the HALLMARK p53 pathway were highly similar to those obtained for the score-based hRPTECTERT1_35 WGCNA module, with values of 6.0 and 5.3 at 16 h, and 0.3 and 0.8 at 48 h, respectively. This similarity indicates that, despite differences in gene composition (10 genes overlap), score-based modeling captures a consistent p53-related response, resulting in comparable BMC estimates when the pathway response can be reliably modelled. Importantly, the inability to model pathway scores at certain timepoints does not necessarily indicate absence of pathway activation but likely reflects limitations of modeling pathway scores.

#### Pathway score-based modeling yields more conservative benchmark concentrations for strong responses

Comparison of score-based BMCs with median-based BMCs revealed a systematic pattern. At the lower end of the BMC distribution, score-based pathway modeling produced lower BMC estimates compared to median-based approaches. At the upper end of the distribution, however, the median-based estimates were relatively lower, indicating that the direction of conservativeness depended on the magnitude of the pathway BMC (Figure 5B**, C**). This behavior was observed consistently in both EG- and NES-based analyses.

Consistent with this pattern, when focusing on the p53 signaling pathway, score-based BMCs were lower than the median-based BMCs (**Table 6**). This suggests that pathways showing strong activation, with many CRGs in the pathway, pathway-level score-based modeling may yield more conservative estimates by capturing the integrated pathway response rather than individual gene effects. At the same time, these findings underscore the absence of a universally “most conservative” approach, as the relative stringency of each method depends on the strength and coherence of the underlying pathway response.

**Table 6.**
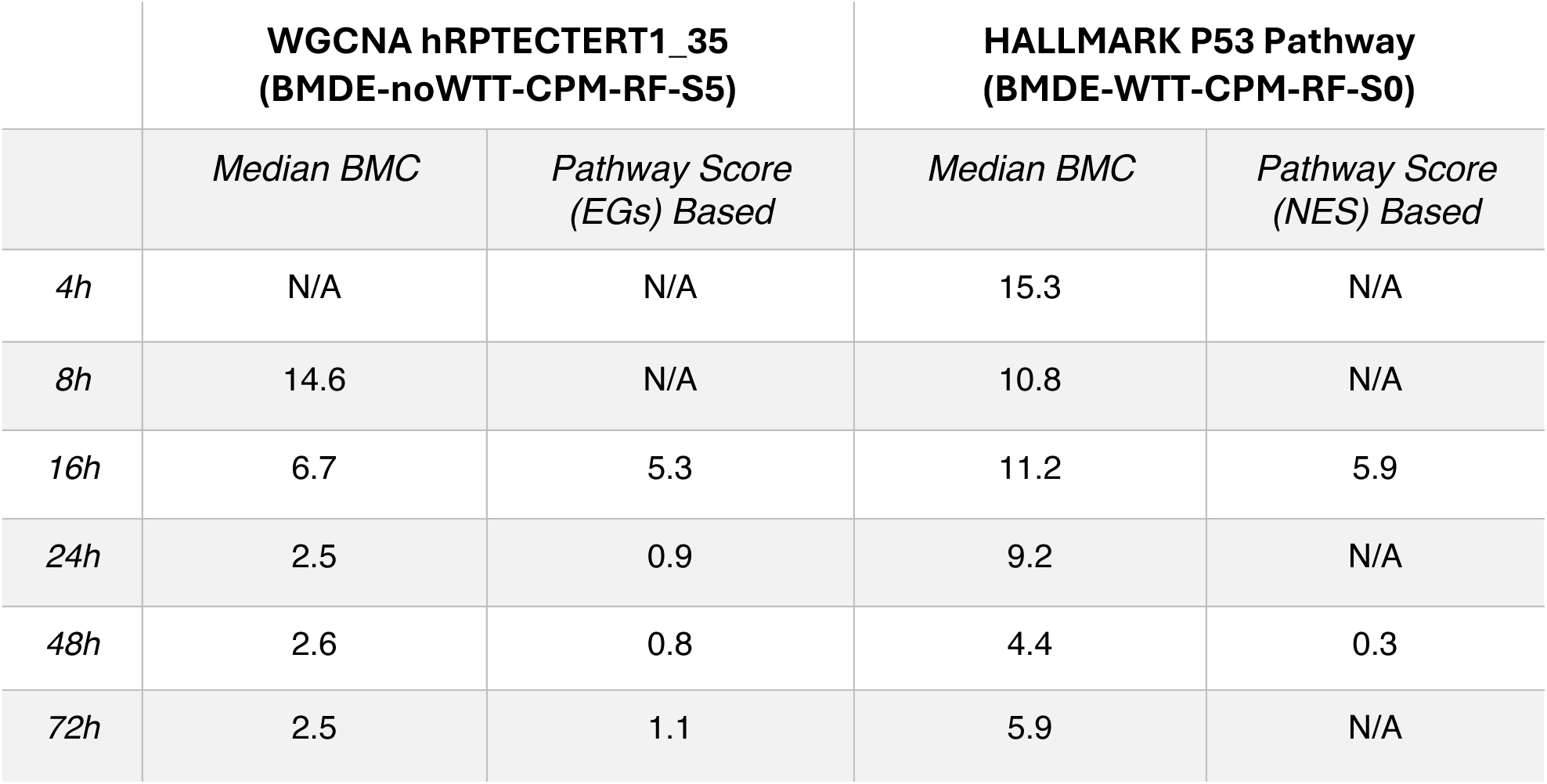
Aggregated median BMCs and pathway score-based BMCs for WGCNA eigengene scores (EGs) and HALLMARK normalized enrichment scores (NES) across timepoints. N/A denotes pathways excluded after post-model filtering or not classified as responsive (e.g. fewer than three responsive genes for BMDE-noWTT & BMDE-WTT, or <5% gene coverage for BMDE-noWTT).

|  | **WGCNA hRPTECTERT1_35**<br>**(BMDE-noWTT-CPM-RF-S5)** |  | **HALLMARK P53 Pathway**<br>**(BMDE-WTT-CPM-RF-S0)** |  |
| --- | --- | --- | --- | --- |
|  | *Median BMC* | *Pathway Score*<br>*(EGs) Based* | *Median BMC* | *Pathway Score*<br>*(NES) Based* |
| 4h | N/A | N/A | 15.3 | N/A |
| 8h | 14.6 | N/A | 10.8 | N/A |
| 16h | 6.7 | 5.3 | 11.2 | 5.9 |
| 24h | 2.5 | 0.9 | 9.2 | N/A |
| 48h | 2.6 | 0.8 | 4.4 | 0.3 |
| 72h | 2.5 | 1.1 | 5.9 | N/A |

As differences between score-based and median-based BMC estimates may reflect how comprehensively a pathway is represented, we next evaluated whether the extent of pathway coverage influenced the differences observed between median BMCs and pathway score-based BMCs, under the hypothesis that greater gene coverage would reduce discrepancies between the two approaches. For the WGCNA modules, pathway coverage was significantly associated with the absolute BMC difference at 24 h (p = 0.013), and 72 h (p = 0.00014), indicating improved agreement between the two approaches when a larger fraction of the module was represented in the median BMC calculation. In contrast, at lower coverage levels (e.g. around 30% at 48 h), substantial variability between the two BMC estimates was observed (Figure 5D).

Taken together, pathway score-based modeling and median aggregation yielded systematically different BMC estimates, with their relative conservativeness depending on gene coverage. Overall, for strongly responding pathways such as the p53 pathways following cisplatin exposure, score-based modeling consistently yielded lower BMC estimates for both NES and EGs. In contrast, median gene-level aggregation was more conservative for pathways with weaker responses.

## Discussion

In this study, we evaluated how methodological choices across independently implemented workflows influence transcriptomic BMC and tPOD derivation. By comparing normalization, modeling, and pathway-level aggregation strategies, we show that methodological variability can substantially affect BMC estimates, with the largest differences arising from the use of different BMD modeling software. However, convergence across workflows increased when transcriptomic responses were stronger, particularly at later timepoints. Gene- and pathway-level analyses captured complementary aspects of transcriptomic sensitivity, while pathway-level results are strongly influenced by pathway definition and gene set size. Together, these findings identify key sources of variability in transcriptomic concentration-response analysis and support the development of best practices for NGRA-relevant applications.

Variation in data normalization and filtering introduced substantial differences in the data prior to BMC modeling, which is in line with previous findings (Haber et al. 2018; Mezencev and Auerbach 2020). This effect is reflected in discrepancies between workflows using the same modeling software but applying different preprocessing strategies. This underscores the importance of streamlined and standardized preprocessing workflows (e.g. R-ODAF) within NGRA frameworks to reduce variability from quality control and normalization steps and to support more consistent interpretation of transcriptomic data for regulatory decision-making (Verheijen et al. 2022).

In addition, prefiltering can lead to large differences in the set of genes retained for modeling, especially at early timepoints when transcriptomic responses are noisy (Webster et al. 2015; Farmahin et al. 2017; Nöth et al. 2025). Interestingly, when prefiltering was not used as an exclusion criterion, gene retention after post-model filtering was highly comparable across workflows, suggesting that post-model criteria alone can help harmonize gene selection. This was particularly evident at later timepoints, where many prefilter-flagged genes passed post-model filters, including goodness-of-fit thresholds such as R^2.^ ≥ 0.8, indicating modellable and potentially biologically relevant concentration-response patterns rather than spurious signals. Such discrepancies are consistent with the behavior of trend-based statistical tests, such as the Williams Trend Test (WTT), which are designed to detect evidence of monotonic concentration-related changes but do not evaluate the shape or quality of fitted concentration-response models (Bretz 2006), as opposed to a quadratic trend test in DRomics, which evaluates linear and non-linear (curvature) trends across concentration levels (Delignette-Muller et al. 2023 Apr 27). As a result, statistically significant yet relatively shallow responses may be excluded by prefiltering despite yielding robust model fits, underscoring the importance of carefully balancing statistical significance thresholds with model-based performance criteria when defining CRGs.

Consistent with the time-dependent stabilization of transcriptomic responses, sensitivity differed by analysis level: 24 h showed the lowest gene-level tPODs, whereas pathway-level aggregation of gene BMCs consistently identified 48 h as the most sensitive timepoint, in line with previous findings (Chauhan et al. 2016). This shift likely reflects progressive coordination of individual gene responses into coherent biological pathways over time, resulting in more integrated and stable pathway-level perturbations. From an NGRA perspective, these findings highlight that the choice of sampling timepoint should be aligned with the intended level of biological interpretation. Gene-level analysis at earlier timepoints (e.g. 24 h) may provide higher sensitivity for detecting molecular initiating or early key events, while later timepoints (e.g. 48 h) may be more informative of pathway-level analyses that support biological interpretation, weight-of-evidence integration, and regulatory decision-making (Kinaret et al. 2020).

Cross-workflows correlations of pathway-level BMCs (HALLMARK) were generally weaker than those observed at the gene level. This reduction likely reflects both aggregation of heterogenous gene responses and the smaller number of pathway-level features, as subsampling gene-level BMCs to a comparable number of data points similarly reduced correlations. In contrast, pathway-level BMCs for the p53 pathway at 48 h converged across workflows within a comparable range, consistent with its well-established role in the mode of action of cisplatin (Rabik and Dolan 2006; Basu and Krishnamurthy 2010; Thompson and Joy 2024). Thus, while pathway aggregation may reduce overall inter-workflow concordance, it can yield consistent estimates when pathway responses are robust.

To reflect a more real-word scenario, we evaluated how pathway definition influences pathway-level BMCs. Comparison of p53-related pathways derived from different sources revealed substantial heterogeneity in gene composition (Bao et al. 2022; Beebe-Wang et al. 2022): no single gene was shared across all five p53 pathway definitions considered, and only one gene (DDB2) overlapped across five of the six pathways (Bao et al. 2022). These findings highlight that pathway-level results are highly dependent on pathway annotation choices, even when representing the same biological process (Mubeen et al. 2019).

To explore whether more focused gene sets could mitigate variability in pathway-level BMCs, we examined the dispersion of gene-level BMCs within different pathway definitions. As expected, larger curated pathways exhibited greater dispersion in individual gene BMCs, reflecting their broad biological scope. This heterogeneity increases the likelihood that only a subset of pathway genes is actively driving the pathway-level response. In contrast, gene-level BMCs within WGCNA-derived modules and curated biomarker gene sets were less dispersed, indicating more coherent transcriptional responses. This aligns with the design of WGCNA modules, which are data-driven and system-specific, as well as the curated biomarker gene sets such as GENOMARK and TGx-DDI that represent defined stress and DNA damage responses (Li et al. 2015; Ates et al. 2018; Costa et al. 2024). Interestingly, gene-level BMC estimates from a core set of p53-associated genes, defined as genes present in at least three of the evaluated p53 pathway definitions, were highly consistent across workflows. This suggests despite variability in broader pathway definitions, a well-curated core gene set can reproducibly capture the central transcriptomic response to cisplatin and yield comparable BMC estimates across analytical workflows. Overall, pathway-level analyses are informative when pathway composition, gene-level coverage, and aggregation criteria are carefully considered. Lenient summarization of large, heterogeneous gene sets (such as HALLMARK, KEGG or WikiPathways) may yield unstable estimates, whereas ensure adequate gene coverage and using complementary approaches such as overrepresentation analysis can improve mechanistic interpretability. Refining aggregation strategies and focusing on biologically coherent core gene sets may further enhance the robustness of pathway-based tPODs for NGRA (Khatri et al. 2012).

In addition to median aggregation of gene-level BMCs, alternative strategies exist that derive BMCs directly from pathway-level activity scores (Harrill, Everett, Haggard, Word, et al. 2024). We therefore compared pathway-score derived BMCs (using HALLMARK-normalized enrichment scores (NES) and eigengene scores (EGs) derived from WGCNA modules), with median-based BMCs from the same pathway sources. Overall, pathway score-based modeling yielded lower BMC estimates at the lower end of the BMC distribution (Harrill, Everett, Haggard, Bundy, et al. 2024), whereas at higher BMC levels the relative conservativeness shifted toward median-based aggregation. It is important to note that failure to derive BMCs from NES- or EG-based modeling does not imply a lack of pathway activation but may instead reflect challenges in fitting a robust concentration-response model. Enrichment-based scores aggregate large gene sets, which can dilute strong signals from a subset of responsive genes (Subramanian et al. 2005). In contrast, median aggregation is highly sensitive to gene selection and pathway coverage, as it relies on the subset of genes passing post-model filters. This sensitivity likely underlies the lower BMC estimates observed at higher BMC ranges, particularly when pathway coverage is limited. For NGRA applications, score-based modeling may therefore be preferable for well-coordinated pathway responses, whereas median aggregation requires sufficient gene-level coverage to avoid unstable or overly sensitive estimates.

Several limitations of this study should be acknowledged. First, the analyses were conducted using a single reference compound, cisplatin, which has a well-characterized mode of action and elicits strong transcriptomic responses for specific pathways of interest (Thompson and Joy 2024). While this study provided a useful benchmark for comparing methodological choices across the different institutions involved, extending the analysis to compounds with less well-defined or more complex modes of action would help evaluate the consistency of conclusions drawn using different analytical approaches. Second, this work is intended to provide methodological guidance and highlight best practices for transcriptomic BMC and pathway-based analyses, rather than to prescribe a single analytical workflow. Many of the observed differences reflect trade-offs between sensitivity, robustness, and interpretability, underscoring the importance of context-specific decision making depending on study objectives and regulatory use cases. Finally, while analytical pipelines were intentionally implemented independently across partners, substantial effort was devoted to harmonizing reporting through a shared BMC framework, specifically developed for this study. This framework offers a practical step toward increased transparency by standardizing how BMC outputs can be summarized, which can facilitate cross-study comparison, internal interpretation, and regulatory evaluation, even when analytical choices differ.

In conclusion, this study highlights how methodological variability profoundly influences the identification of tPODs for NGRA of chemicals, which underscores the need for a higher level of methodological harmonization to ensure consistency in the increasing use of omics data in regulatory contexts. Although omics technologies offer unprecedented resolutions to characterize molecular perturbations, our comparison of five independent analysis workflows shows that even when using the same experimental dataset, differences in preprocessing choices, BMD modeling tools, pathway selection, and tPOD derivation methods can lead to markedly different downstream results, emphasizing the need for higher level methodological harmonization.

## Acknowledgement

This research was funded by the European Union’s Horizon Europe Partnership for the Assessment of Risks for Chemicals (PARC) (grant agreement no. 101057014); the IMI TransQST project (grant agreement no. 116030): and the European Union’s Horizon 2020 RISK-HUNT3R Initiative (grant agreement no. 964537).

