## Supplemental Figure 1 for "Analytical Choices Drive Toxicogenomic Potency Estimates: A Systematic Evaluation of Transcriptomic Points of Departure"

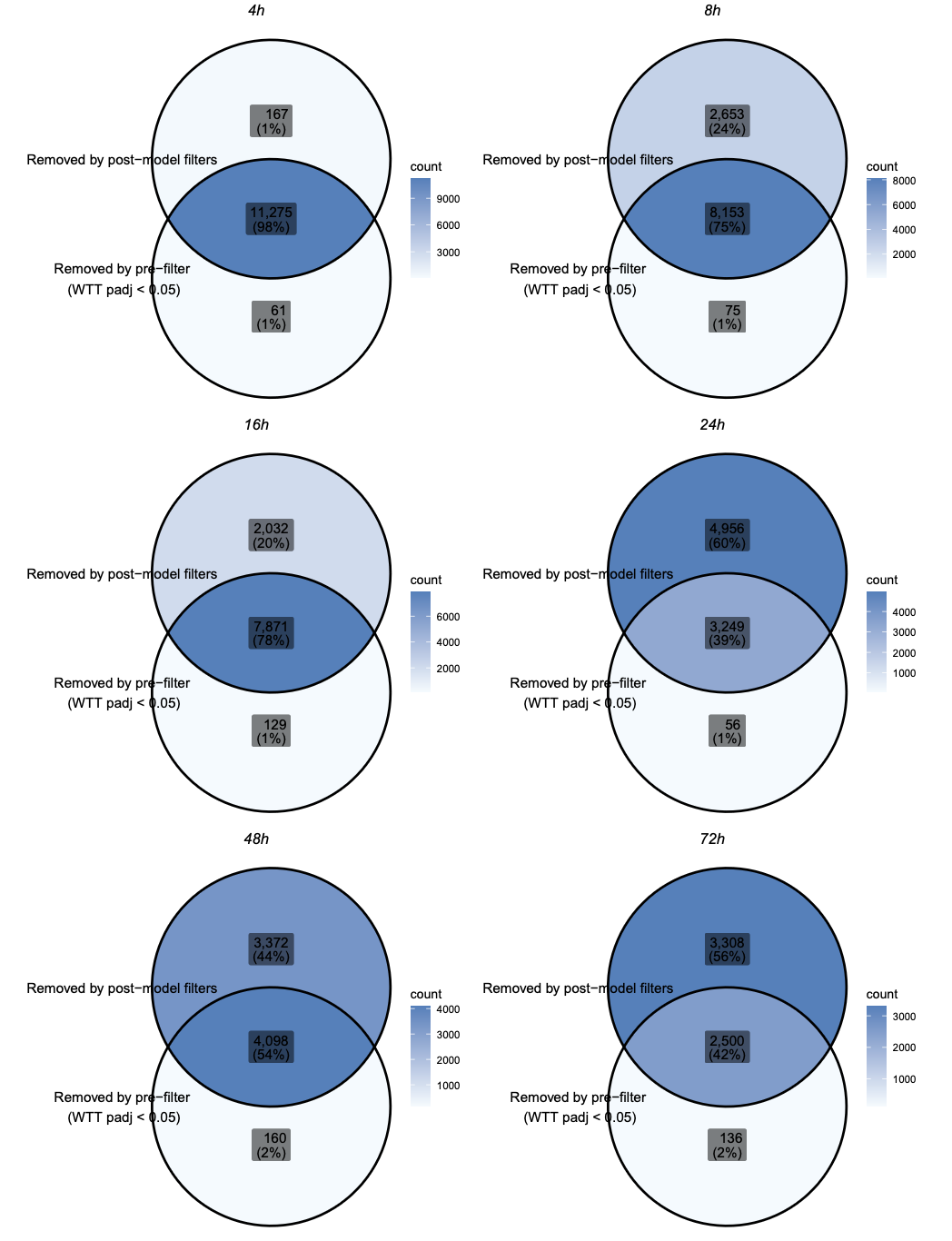

**Supplemental Figure 1.** Comparison of genes excluded by prefiltering (Williams Trend Test, adjusted p-value < 0.05), and genes removed by post-model filtering (BMC >= 0.01, BMC <= 50, BMD/BMCL <= 20, BMCU/BMC <= 20, R-squared >= 0.8). Results are shown for the BMDE-noWTT-CPM-RF-S5 pipeline.
