## Supplemental Figure 2 for "Analytical Choices Drive Toxicogenomic Potency Estimates: A Systematic Evaluation of Transcriptomic Points of Departure"

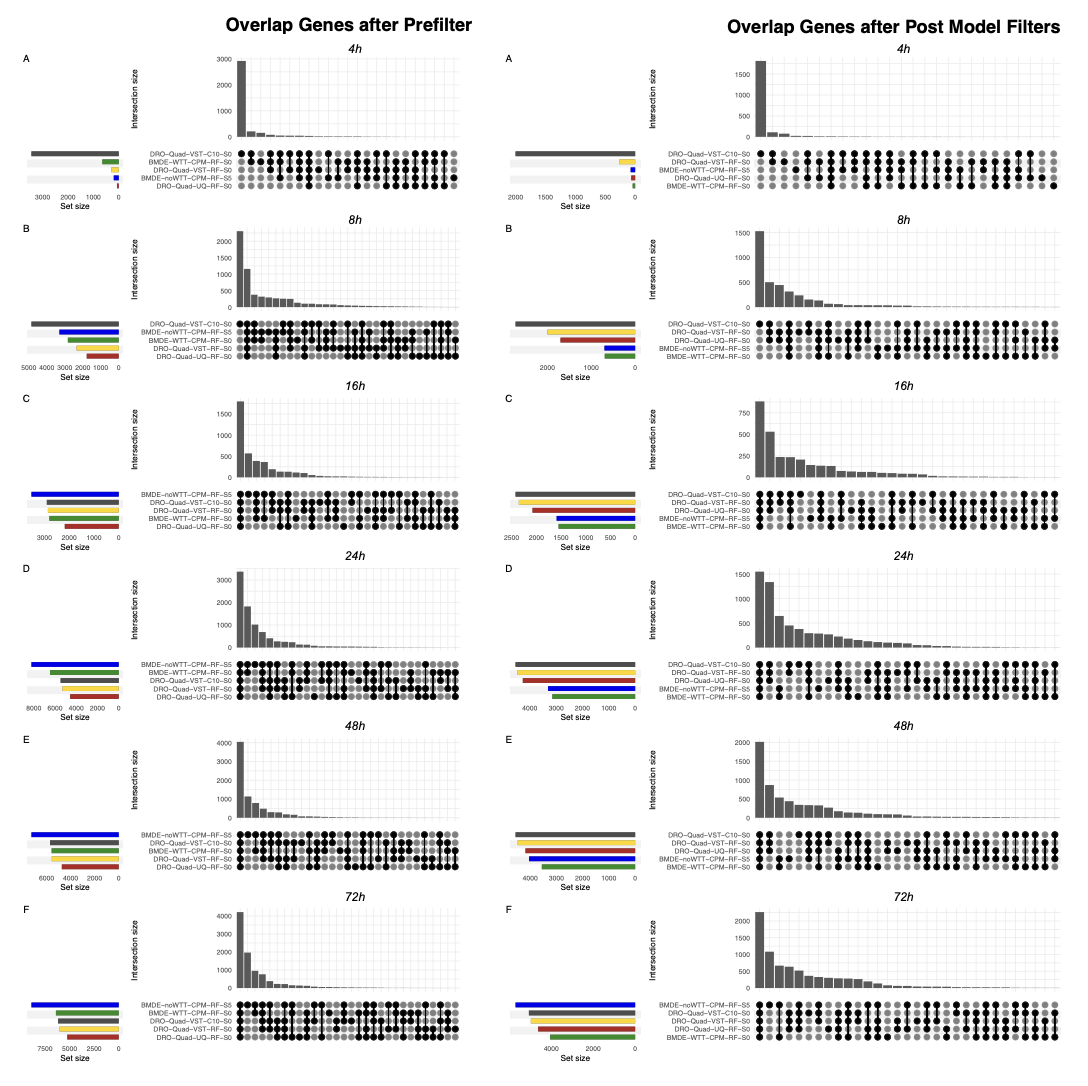
**Supplemental figure 2.** Overlap of genes retained across analytical methods following prefiltering (left) and post-model filters (right). Colors represent different methods.
