## Supplemental Figure 3 for "Analytical Choices Drive Toxicogenomic Potency Estimates: A Systematic Evaluation of Transcriptomic Points of Departure"

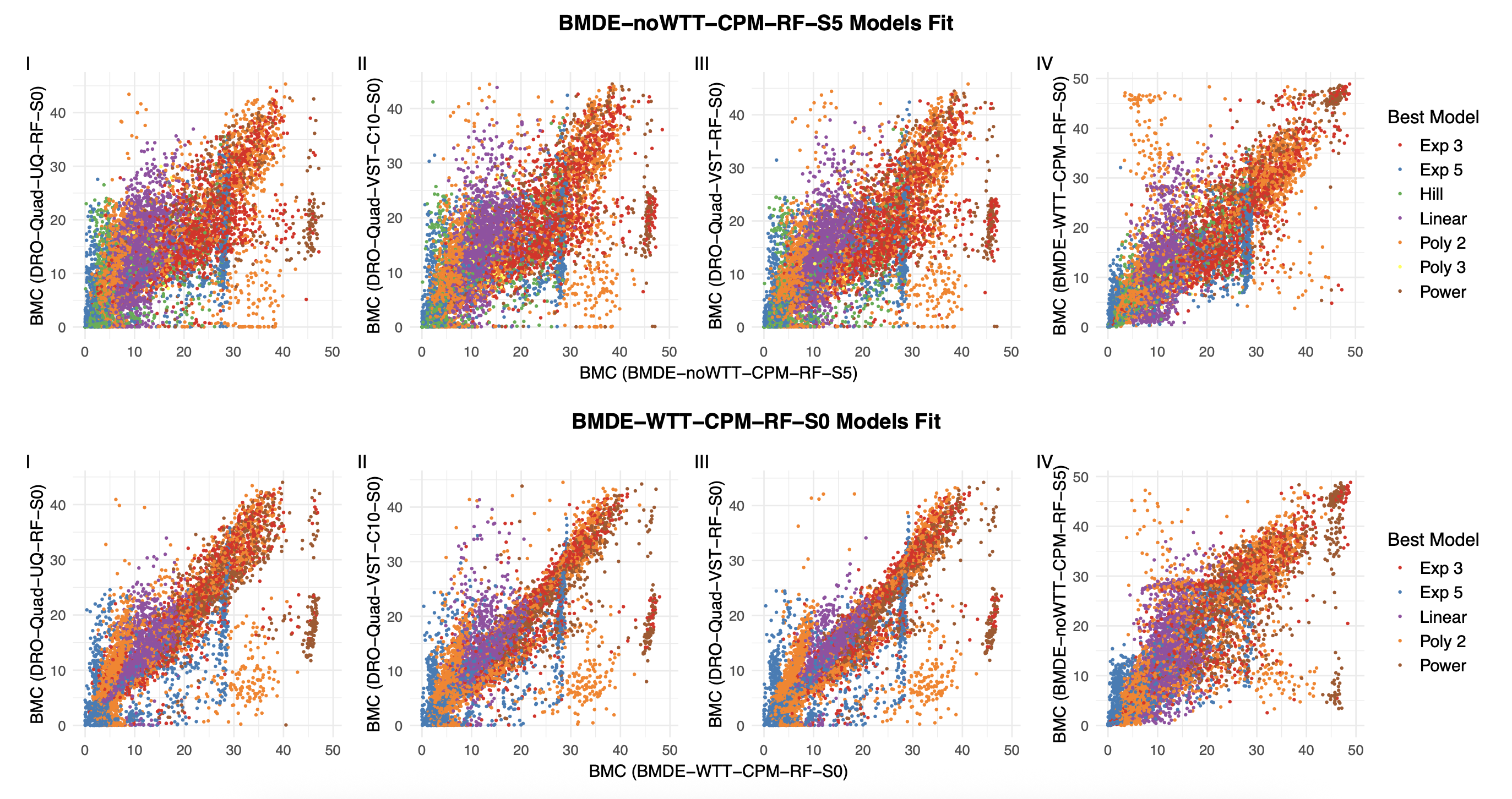

**Supplemental Figure 3.** Correlation plots of best-fit models within the BMDExpress workflows (BMDE-noWTT-CPM-RF-S5 & BMDE-WTT-CPM-RF-S0), illustrating model-specific patterns contributing to the accumulation of benchmark concentrations (BMCs).
