## Supplemental Figure 4 for "Analytical Choices Drive Toxicogenomic Potency Estimates: A Systematic Evaluation of Transcriptomic Points of Departure"

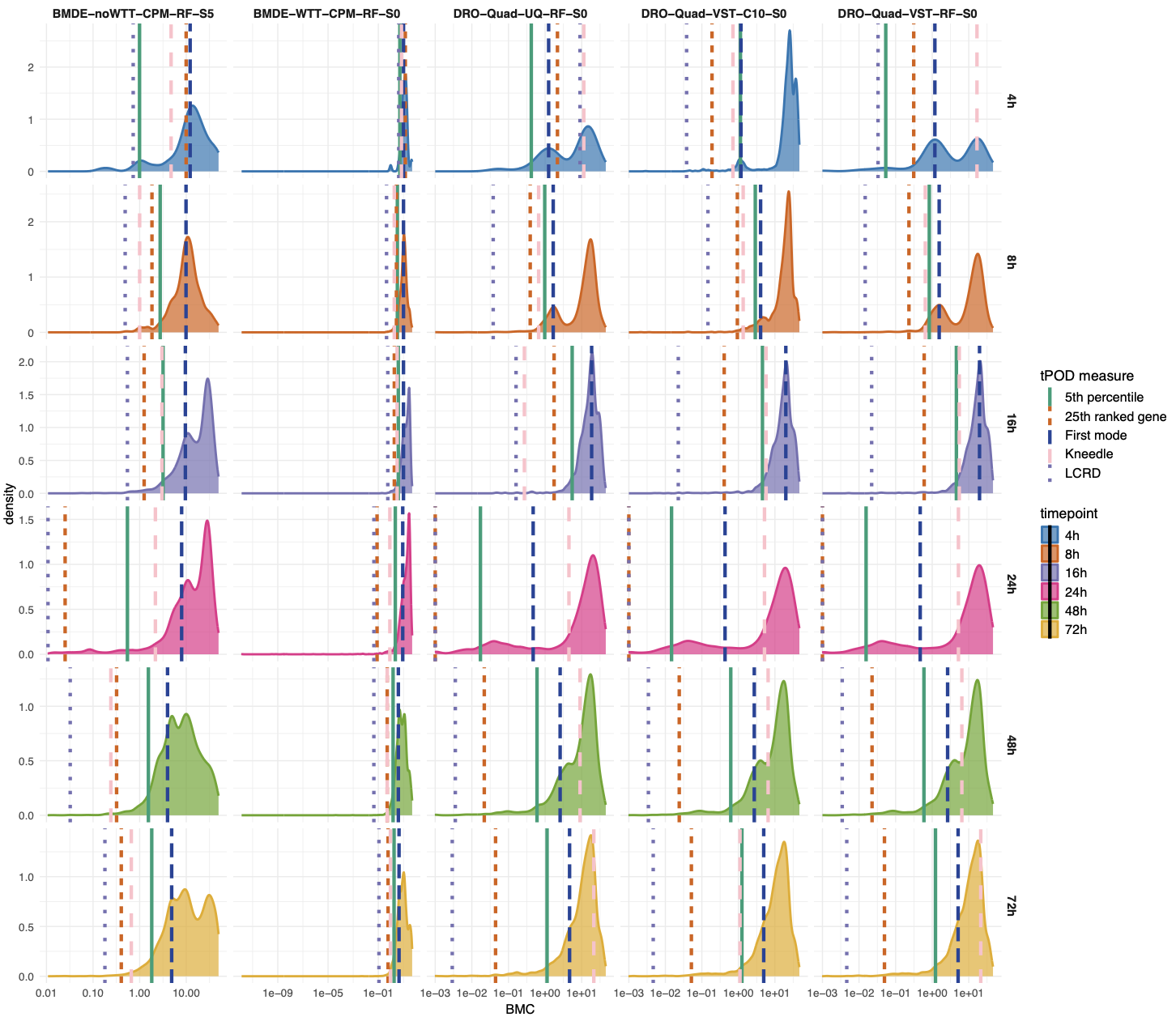

**Supplemental Figure 4. Distribution of gene-level BMCs with corresponding** tPOD **estimates.** Vertical lines indicate the evaluated tPOD metrics (5^th^ percentile, 25^th^ ranked gene, first mode of the distribution, inflection point [Kneedle] and LCRD), with different line types and colors distinguishing the metrics. Rows represent the different timepoints.
