## Supplemental Methods for "Analytical Choices Drive Toxicogenomic Potency Estimates: A Systematic Evaluation of Transcriptomic Points of Departure"

### **BMDE-noWTT-CPM-RF-S5 workflow**

#### *TempO-Seq data preprocessing*

TempO-Seq analysis was performed using a whole transcriptome probe panel, and raw read counts were provided by BioClavis. All subsequent analyses were performed in R (version 4.5.0). To ensure data quality, several preprocessing steps were applied. First, probe-to-gene mapping was performed using the Human Whole Transcriptome 1.2 manifest (realigned in 2023). Next, a relevance filter adapted from the R-ODAF pipeline was applied, resulting in the removal of 8477 probes (Verheijen et al. 2022). Probe counts mapping to the same gene were aggregated, yielding 11531 unique genes for downstream analysis.

Samples with a total library size below 500,000 reads were excluded, leading to the removal of one sample. To assess replicate quality, pairwise correlations were calculated among replicates within each treatment group. For each group, a correlation threshold was defined as the maximum pairwise correlation minus three times the interquartile range (3*IQR). Replicates with a maximum pairwise correlation below this threshold were excluded. This step resulted in the removal of three additional samples, for a total of four samples excluded from further analysis. Finally, raw counts were normalized to counts per million (CPM), followed by a log2 transformation after adding a pseudo count of 1 to avoid infinite values, enabling comparison across samples.

#### *Differential Expression Analysis*

Differential expression analysis was conducted using DESeq2 (version 1.48.2) (Love et al. 2014). Departing from the standard DESeq2 workflow, which typically estimates a single log2 fold change (log2FC) per treatment condition by aggregating replicates, we adapted the analysis to calculate log2FC values for each replicate individually. For this, each treatment replicate was compared against the pooled control replicates (n=6) at the corresponding timepoint. This approach was necessary to generate replicate-level log2FC values required as input for downstream analysis in BMDExpress3.

#### *TXG-MAPr module projection*

To assess biological responses at the pathway level, log2FC values were uploaded in the TXG-MAPr platform (Callegaro et al. 2021). This tool enables the projection of new gene expression data onto a precomputed Weighted Gene Co-Expression Network Analysis (WGCNA) map (Langfelder and Horvath 2008). In our case, the map was previously generated in-house using a large RPTEC-TERT1 dataset, which included transcriptomic profiles from exposures to over 50 compounds across multiple concentrations and two timepoints (van Kessel et al. 2025 Nov 20). The resulting network consists of 291 gene co-expression modules representing diverse biological processes. By projecting our log2FC values onto this reference map, we obtained eigengene scores summarized expression profiles for each module in every sample, facilitating interpretation of the underlying biology in a structured pathway-level context.

#### *Benchmark Concentration Analysis*

##### Gene level analysis

Benchmark concentration modeling was conducted using BMDExpress3 (version 3.20) on normalized expression values (log2CPM), log2FC values and eigengene scores as described above (Yang et al. 2007). No prefiltering was applied, as we aimed to retain all genes for modeling without exclusion based on preliminary statistical tests such as the Williams Trend Test (WTT). The BMC analysis utilized the suite of EPA BMDS models implemented in BMDExpress3, including Hill, Power, Linear, Polynomial (2^nd^ and 3^rd^ order), and Exponential (3^rd^ and 5^th^ order) models. Benchmark response (BMR) thresholds were defined using the standard deviation method with a BMR factor of 1.021 (5%).

Post-modeling, genes with benchmark concentrations (BMCs) below 10% of the lowest tested concentration (0.01 µM) or above the highest tested concentration were excluded. To ensure reliable concentration-response estimates, genes were further filtered based on model uncertainty: those with a BMCL/BMC or BMC/BMCU ratio >= 20 were removed. Finally, genes with a model fit R^2^ < 0.8 were excluded to ensure adequate model performance.

##### Functional classification

All concentration-responsive genes that passed the previously described quality filters were assigned to gene sets, provided that at least three such genes were present within the set, and 5% of the gene set is covered by those genes. For functional annotation and cross-workflow comparison, we used the MSigDB HALLMARK gene sets (2023) (Liberzon et al. 2015). Additionally, WGCNA-derived RPTEC-TERT1 modules were also treated as gene sets to obtain a BMC value per module (van Kessel et al. 2025 Nov 20). Pathway-level BMCs were calculated by taking the median BMC of all qualifying genes within each gene set.

### **BMDE-WTT-CPM-RF-S0 workflow**

This workflow follows an approach that has been implemented in previous studies (Thienpont et al. 2024; Thienpont et al. 2025). Initially, this workflow was developed to derive transcriptomic Points of Departures (tPODs) for two biomarkers of genotoxicity called GENOMARK and TGx-DDI. Given that Cisplatin is a known genotoxicant, we found it relevant to include these two biomarkers in our analysis. Thus, choices in our workflow have been influenced by the analysis of genotoxic biomarkers.

#### *Transcriptomics data preprocessing*

First, probe-to-gene mapping was performed using the Human Whole Transcriptome 1.2 manifest (realigned in 2023). The relevance filter, adapted from the R-ODAF pipeline (Verheijen et al. 2022), was then applied, resulting in the removal of 8,477 probes. Probe counts mapping to the same gene were aggregated, yielding 11,531 unique genes for downstream analysis.

The resulting dataset was normalized using the DESeq2 R package. Normalized counts were log2-transformed for downstream analyses.

#### *Benchmark Concentration Analysis*

##### At the gene level

To conduct the BMC analysis, log₂-normalized read counts were uploaded into BMDExpress v3, following the recommendations outlined in the U.S. NTP Approach to Genomic Dose-Response Modeling report (NTP 2018; Phillips et al. 2019). Gene expression data were prefiltered using the Williams trend test, retaining genes with a permutation p-value < 0.01 (based on 500 permutations) and an absolute linear fold change (FC) > 1.5 for at least one concentration relative to matched controls. For genes passing these filters, concentration-response modeling was performed using following EPA BMDS models: Linear, Exponential (3 and 5), Polynomial (2nd degree), and restricted Power (exponent ≥ 1). In BMDExpress v3, the optimal polynomial degree was determined using a nested chi-square test, after which the Akaike Information Criterion (AIC) was applied to select the best-fitting model, balancing goodness-of-fit with model complexity. The benchmark response (BMR) was set to 1 SD, following the recommendations of the U.S. NTP expert panel. BMC estimates were further filtered using quality criteria: only models with a goodness-of-fit p-value > 0.1, a BMC/BMCL ratio < 20, a BMCU/BMCL ratio < 40, and BMC values below the highest tested concentration (50 µM) were retained. Here, BMCL and BMCU represent the lower and upper confidence limits of the BMC, respectively.

##### At the biological entity level

The median of gene set BMCs were used to derive tPODs for the hallmarks and genotoxic biomarkers GENOMARK and TGx-DDI.

### **DRO-Quad-UQ-RF-S0 workflow**

#### *Raw Data Processing, Normalisation, and Filtering*

This workflow was developed for analyzing BioSpyder TempoSeq count data, which has already undergone manufacturer platform-specific preprocessing steps (library preparation, base calling, raw data filtering, sequence alignment, gene quantification). Count data were preprocessed using R v4.3.3 (*limma* v3.56.2, *edgeR* v3.42.4, *dplyr* v1.1.4, and *baseR*) (Robinson et al. 2009; Ritchie et al. 2015). Quality control visualizations were used to assess the efficacy of preprocessing steps. Primarily, library sizes were calculated, and a threshold of 500,000 read counts was set. All samples passed library size cutoffs. Next, normalisation methods were assessed for applicability and included no normalisation (none), upper quartile (UQ), trimmed mean of M-values (TMM), TMM with singleton paring (TMMwsp), and relative log expression (RLE) [1]. After assessment, the UQ normalisation method was applied (Li et al. 2020). A relevance filter was applied as per Verheijen *et al.* (Verheijen et al. 2022). Next, probes were mapped to the corresponding EntrezIDs and those corresponding to the same gene were summed. Correlation analysis showed improved correlation values after preprocessing and overall good accordance between controls.

#### *Concentration-Response Modeling and Statistical Analysis*

BMC analysis was carried out using the *DRomics* v2.5-2 package in R to derive tPODs, using the quadratic trend test (Larras et al. 2018), with multiple testing correction using the Benjamini-Hochberg methods and a false discovery rate (FDR) threshold of 0.01 for concentration-responsive genes, and confidence interval calculations using 1,000 bootstrap iterations. A parametric modeling approach is performed by *DRomics*, whereby multiple models are fitted (linear, Hill, exponential, Gauss-probit, and log-Gauss-probit functions). This allows for the assessment of monotonicity and biphasic trends. The best fit model is selected based on the second order Akaike Information Criterion (AICc). Benchmark response was defined as 1 standard deviation (1 SD) from the control mean (zSD). A maximum of 1,000 iterations were allowed for model fitting, and a 95% confidence level with a tolerance of 0.5 was applied to derive BMC intervals. Items for which bootstrap confidence intervals could not be computed were excluded from downstream analysis.

#### *Gene and Pathway Level tPOD Derivation*

The R packages *KEGGrest* v1.40.1 and *clusterProfiler* v4.8.3 were used for pathway enrichment analysis using KEGG (2021) and MSigDB Hallmark (2021) databases (Kanehisa and Goto 2000; Liberzon et al. 2015). Direct output from the BMC analysis generated gene-level BMC values. In addition, gene set/pathway-level BMC estimates were performed as per the U.S. NTP recommendations (National Toxicology Program 2018). For features to be included in gene set analysis, the following criteria were implemented for the best fit models: (1) must demonstrate convergent BMC, BMCL, and BMCU values; (2) the BMC must be less than the highest positive concentration used in the study; (3) it should not map to more than one gene; (4) the BMCU-to-BMCL ratio should be less than 40. Concentration-responsive items were filtered based on these criteria, and genes were parsed into sets that met the following conditions: contain at least 3 genes and populate at least 5% of the total annotated gene number.

### **DRO-Quad-VST-RF-S0 workflow**

R version 4.4.2 was used as the general platform for analyses. First, a quality control of the samples was done based on sequencing depth. A histogram of sequencing depths was made. No clear outliers were observed, and all samples had a sequencing depth larger than 500,000, so no samples were removed. Then, the data and metadata were loaded in DESeq2 (version 1.44.0). A factor with every unique combination of concentration and time was used to set up the design matrix. Afterwards, a relevance filter was applied to remove genes that had low expression and were not homogenously detected within their condition/group. More specifically, all genes that did not have at least 75% of their replicates expressed at 1 CPM in any one of the experimental conditions were excluded. A correlation matrix was made based on variance stabilized transformation (vst) result of the read counts. The replicate correlation was always higher than 80%, so no samples were removed. Finally, raw counts whose probes were mapped to the same entrez id were summed.

Concentration–response characterization was performed using **DRomics** (version 2.6.2) (Larras et al. 2018). RNA-seq count data were imported into DRomics with the function RNAseqdata(). As recommended, raw counts were used, and variance stabilization was applied with the *vst* option (variance-stabilizing transformation), which is preferred over *rlog* when the number of samples exceeds 30. This transformation automatically normalizes for library size and applies a log2 scale. Significantly responding genes were identified with the function itemselect, using a quadratic trend test with a false discovery rate (FDR) threshold of 0.01. Concentration–response modeling was then carried out by fitting five families of models designed to capture monotonic and biphasic responses: Hill, linear, exponential, Gauss-probit, and log-Gauss-probit. For each gene, the best-fitting model was selected based on the second-order Akaike Information Criterion (AICc). Benchmark concentration (BMC) values were estimated with the function bmdcalc. Confidence intervals for BMCs were obtained by bootstrapping with 1,000 iterations. Only genes with finite BMC point estimates and bounded confidence intervals were retained for downstream analyses.

The hallmark gene sets form the molecular signature database were loaded in r with the function msigdbr(species = “Homo sapiens”, category = “H”) from the package “msigdbr” (version 7.5.1). The median and IQR for each of the gene sets were calculated, and only gene sets with at least 3 genes were retained.

### **DRO-Quad-VST-C10-S0 workflow**

#### *Transcriptomics data preprocessing*

All analyses were performed in R (version 4.4.2). Probe-to-gene mapping was conducted using the Human Whole Transcriptome 1.2 manifest (realigned in 2023). When multiple probes mapped to the same Entrez Gene ID, their counts were summed to obtain gene-level expression values. Prior to concentration–response analysis, a minimal expression filter was applied. Genes were retained if they had at least 10 raw counts in a minimum of three samples. No additional CPM-based relevance filtering was applied for the BMC modelling step.

Sequencing depth and sample relationships were assessed descriptively. All samples passed quality control and were retained for downstream analyses. Variance-stabilizing transformation (VST) from DESeq2 was used for exploratory analyses (e.g. PCA), but not for concentration–response modelling.

#### *Benchmark Concentration Analysis*

Concentration–response analysis was carried out using the DRomics package (version 2.5-2) in R (Larras et al. 2018). Raw gene-level counts were imported into DRomics by implementing the *formatdata4DRomics* and *RNAseqdata* functions. Internal normalization and transformation were performed within DRomics prior to modelling. Concentration-responsive genes were identified using the quadratic trend test, applying a Benjamini–Hochberg false discovery rate (FDR) threshold of 0.01. Parametric concentration–response models were fitted using the linear, Hill, exponential, Gauss–probit and log-Gauss–probit model families. For each gene, the best-fitting model was selected based on the second-order Akaike Information Criterion (AICc). Benchmark concentration (BMC) values were derived using the BMR-zSD approach, defining the benchmark response as a 1 standard deviation change from the control mean. Confidence intervals were estimated by bootstrap with 1000 iterations. Only genes with finite BMC confidence intervals were retained for subsequent analyses. No additional filtering based on BMC magnitude or BMC confidence interval width was applied.

#### *Gene set and pathway-level analysis*

Over-representation analysis was performed separately for each timepoint using the lists of concentration-responsive genes with finite BMC confidence intervals. Enrichment analysis was conducted using Reactome (ReactomePA), KEGG, MSigDB Hallmark gene sets (msigdbr), and WikiPathways (clusterProfiler). Default significance thresholds implemented in these tools were applied (Benjamini–Hochberg adjusted p-values). For pathway-level BMC estimation, gene-level BMC values were mapped to gene sets and the median BMC (together with the corresponding median BMCL and BMCU) was calculated for each biological set.
