## Supplemental Table 1 for "Analytical Choices Drive Toxicogenomic Potency Estimates: A Systematic Evaluation of Transcriptomic Points of Departure"

### **Supplemental Tables**

**Supplemental Table 1. Selected variables from the OECD Omics Reporting Framework (OORF).** This table presents a selection of key OORF variables used to compare the analytical workflows across partners. It serves as the main reference document for identifying methodological differences between workflows.

| Description | | DRO-Quad-UQ-RF-S0 | DRO-Quad-VST-RF-S0 | DRO-Quad-VST-C10-S0 | BMDE-noWTT-CPM-RF-S5 | BMDE-WTT-CPM-RF-S0 |
| --- | --- | --- | --- | --- | --- | --- |
| 3.2.3 Analysis of Raw Data | 3.2.3.3.i Raw Data filtering and trimming: Minimum read count | 500 000 | Histogram; no outliers | 500,000 | 500,000 (after probe QC), removed 1 sample | 500,000 |
|  | 3.2.3.5.iv Gene quantification: Description of method for summarizing transcript or probe counts to gene counts | Probes mapped to the same Entrez ID_id were summed | Probes mapped to the same Entrez ID were summed | Probes mapped to the same Entrez ID were summed | Probes mapped to the same Entrez ID were summed | Probes mapped to the same Entrez ID were summed |
| 3.2.4. Data Normalisation | 3.2.4.1.a Normalisation of the Raw Count method | Upper Quartile normalization | Log2 VST | Log2 VST | Log2 CPM | Log2 CPM |
| 3.2.5. Post-Normalisation Data Filtering | 3.2.5.1.a&b Identification and Removal of Low Quality or Outlying Data Sets | PCA + correlation analysis to visualize, no samples removed | Replicate correlation < 0.8, No samples were removed | NA | Replicate correlation - calculate pairwise correlations for all replicates in a treatment condition, find max correlation (X), remove samples if the max pairwise correlation of one replicate within the treatment condition is < X - 3*IQR. This removed 4 samples | Replicate correlation < 0.8: no samples were removed |
|  | 3.2.5.2.a Identification and Removal of Low Quality or Outlying Data Sets: Low read count filtering step | Relevance filter | Relevance filter | Minimum raw-count criterion (≥10 counts in at least 3 samples) | Relevance filter | Relevance filter |
| 4.1.5. Statistical Analysis to Identify Differentially Abundant Molecules | 4.1.5.1. Statistical Approach | N/A | NA | N/A | Deseq2 (+/-) | Deseq2 (+/-) |
| 4.3. Data Analysis Reporting Module (DARM) for Benchmark Dose Analysis and Quantification of Biological Potency. | 4.3.2.1. Software | DRomics | DRomics | DRomics 2.5-2 | BMDExpress3 (Version: BMDExpress 3.20.0095 BETA) | BMDExpress3 (Version: BMDExpress 3.20.0095 BETA) |
|  | 4.3.4.1. Statistical Test Performed to Identify Dose-Responsive MSD | Quadratic (Larras et al., 2018) | Quadratic | Quadratic | Williams Trend Test (but not used as pre-filter for BMD input) | Williams Trend test |
|  | 4.3.4.2. Statistical Threshold Applied | FDR < 0.01 | FDR < 0,01 | FDR < 0.01 | p-value < 0.05 (no exclusion critérium) | p-value < 0.01 & fold filter >= 1.5 |
|  | 4.3.4.3. Statistical Multiple Testing Correction Method | Benjamini-Hochberg | Benjamini-Hochberg | Benjamini-Hochberg | N/A | N/A |
|  | 4.3.4.4. Additional Statistical Test Parameters | Bootstrap confidence intervals = 1000 | Number of bootstrap to calculate confidence interval = 1000 | 1000 samples by bootstrap | *Number of permutations: 10000* | permutations: 500 |
|  | 4.3.4.5. Additional Filtering | NA | NA | NA | N/A | N/A |
|  | 4.3.5.1. BMD Modeling Approach | Parametric | Parametric | Parametric | Parametric | Parametric |
|  | 4.3.5.2. Method for Final Estimation of MSD BMD | best model selection | best model selection | best model selection | best model selection | best model selection |
|  | 4.3.5.3. List of Models Fit to the Data | Linear, Hill, exponential, Gauss-probit and log-Gauss-probit (see Larras et al. 2018 for their definition) | Linear, Hill, exponential, Gauss-probit and log-Gauss-probit | linear, Hill, exponential, Gauss-probit and log-Gauss-probit | EPA BMDS models: hill, power, linear, poly 2, poly 3, exponential 3, exponential 5 | EPA BMDS models: power, linear, poly 2, exponential 3, exponential 5 |
|  | 4.3.5.4. Model Averaging Approach | NA | N/A | N/A | *N/A* | N/A |
|  | 4.3.5.5. Method for Determining Benchmark Response (BMR; a.k.a. BMR Type) | BMR-zSD = y0 +/- z*SD, where y0 is the mean control response, and SD is the residual standard deviation of the considered CRC and z is the factor of SD | BMR-zSD = y0 +/- z*SD, where y0 is the mean control response, and SD is the residual standard deviation of the considered CRC and z is the factor of SD | BMR-zSD = y0 +/- z*SD, where y0 is the mean control response, and SD is the residual standard deviation of the considered CRC and z is the factor of SD | Standard Deviation | Standard Deviation |
|  | 4.3.5.6. Benchmark Response (BMR, a.k.a. BMR Factor) | z = 1, so 1SD | z = 1, so 1SD | z = 1, so 1SD | 1.021 (5%) | 1 SD |
|  | 4.3.5.7. Number of Fitting Iterations | 1000 | Bootstrap: 1000 | 1000 | 250 | 250 |
|  | 4.3.5.8. Confidence Interval of the BMD (a.k.a. Confidence Level) | 0.95 with tolerance level of 0.5 | 0.95 | 0.95 | 0.95 | 0.95 |
|  | 4.3.5.9. Power Parameter Restriction | N/A | N/A | Not explicit. | *>=1* | *>=1* |
|  | 4.3.5.10. Variance Assumption | *None* | *None* | None | Constant | Constant |
|  | 4.3.5.11. Model Selection | Best AICc | Best AICc | Best AICc | *Lowest AIC; models were only considered if (1) BMD/BMDL < 20, (2) BMDU/BMD < 20, (3) RSquared >= 0.8, (4) BMD between 1/10 of lowest dose and highest dose.* | Lowest AIC; models were only considered if (1) BMD <= Highest concentration, (2) BMD/BMDL < 20, (3) BMDU/BMDL < 40, (4) RSquared >= 0.8 |
|  | 4.3.5.12. Criteria to Identify Models that Extrapolate Outside the Dose Range | Finite BMD point estimates and CI bounds | Finite BMD point estimates and CI bounds | Finite BMD point estimates and CI bounds | Model was only considered when the Best BMD was between 1/10 of the lowest dose and highest concentration. | Model was only considered when BMD was lower than the highest concentration |
|  | 4.3.6.1. Biological Entity or Biological Set Annotation Used for the Analysis | Human KEGG (2021), MSigDB (Hallmark, 2021) | Hallmark gene sets from MSigDB (*msigdbr* package v25.1.1) | Hallmark (msigdbr package v25.1.1), WikiPathways (clusterProfiler 4.16.0) | RPTEC-TERT1 TXG-MAPr modules, MSigDB (Hallmark, 2023) | GENOMARK, TGxDDI biomarker sets, Hallmark (version 2023.2) |
|  | 4.3.6.2. MSD Acceptance Criteria for Use in Biological Entity of Set Analysis | (1) BMD <= 50; (2) BMDU/BMDL <= 40. | N/A | *N/A* | *N/A* | N/A |
|  | 4.3.6.3. Criteria for Identification of “Active” Biological Sets | Contain at least 3 genes and populate at least 5% of the total annotated gene number. | Gene sets that contained at least 3 genes were retained | Gene sets with an ORA padj < 0.1 were included | At least 3 genes passing all filters, >=5% coverage | GENOMARK and TGxDDI: positive predictions in at least one concentration of any time point.  HALLMARKS: NES associated padj < 0.05 |
|  | 4.3.6.4. Method of Estimating the BMD, BMDL, and BMDU of the Individual Biological Entity or Biological Sets | Median | Median | Median | Median | Median |
