## Supplemental Table 2 for "Analytical Choices Drive Toxicogenomic Potency Estimates: A Systematic Evaluation of Transcriptomic Points of Departure"

**Supplemental Table 2. Benchmark concentrations for the top 15 most sensitive HALLMARK pathways for each workflow at 48 h.** Pathways are ranked by increasing BMC, highlighting the most sensitive pathway-level responses across workflows. The HALLMARK p53 pathway is underlined.

|  | BMDE-noWTT-CPM-RF-S5 | | BMDE-WTT-CPM-RF-S0 | | DRO-Quad-UQ-RF-S0 | | DRO-Quad-VST-C10-S0 | | DRO-Quad-VST-RF-S0 | |
| --- | --- | --- | --- | --- | --- | --- | --- | --- | --- | --- |
|  | **Hallmark pathway** | **BMC** | **Hallmark pathway** | **BMC** | **Hallmark pathway** | **BMC** | **Hallmark pathway** | **BMC** | **Hallmark pathway** | **BMC** |
| 1 | HALLMARK_P53_PATHWAY | 6.89 | HALLMARK_P53_PATHWAY | 9.21 | HALLMARK_E2F_TARGETS | 3.76 | HALLMARK_E2F_TARGETS | 2.36 | HALLMARK_E2F_TARGETS | 2.36 |
| 2 | HALLMARK_SPERMATOGENESIS | 8.38 | HALLMARK_TNFA_SIGNALING_VIA_NFKB | 9.42 | HALLMARK_G2M_CHECKPOINT | 8.32 | HALLMARK_G2M_CHECKPOINT | 4.17 | HALLMARK_G2M_CHECKPOINT | 4.79 |
| 3 | HALLMARK_APOPTOSIS | 9.20 | HALLMARK_ESTROGEN_RESPONSE_LATE | 9.53 | HALLMARK_TNFA_SIGNALING_VIA_NFKB | 9.65 | HALLMARK_HYPOXIA | 4.82 | HALLMARK_HYPOXIA | 6.81 |
| 4 | HALLMARK_TNFA_SIGNALING_VIA_NFKB | 9.22 | HALLMARK_APOPTOSIS | 9.77 | HALLMARK_EPITHELIAL_MESENCHYMAL_TRANSITION | 9.97 | HALLMARK_APICAL_JUNCTION | 6.77 | HALLMARK_MITOTIC_SPINDLE | 7.33 |
| 5 | HALLMARK_EPITHELIAL_MESENCHYMAL_TRANSITION | 9.35 | HALLMARK_EPITHELIAL_MESENCHYMAL_TRANSITION | 9.93 | HALLMARK_INFLAMMATORY_RESPONSE | 10.41 | HALLMARK_GLYCOLYSIS | 6.89 | HALLMARK_KRAS_SIGNALING_DN | 7.73 |
| 6 | HALLMARK_PANCREAS_BETA_CELLS | 9.56 | HALLMARK_PI3K_AKT_MTOR_SIGNALING | 9.97 | HALLMARK_HYPOXIA | 10.70 | HALLMARK_ANGIOGENESIS | 7.06 | HALLMARK_APICAL_JUNCTION | 7.74 |
| 7 | HALLMARK_PI3K_AKT_MTOR_SIGNALING | 9.56 | HALLMARK_PEROXISOME | 10.31 | HALLMARK_COAGULATION | 10.77 | HALLMARK_MITOTIC_SPINDLE | 7.31 | HALLMARK_GLYCOLYSIS | 7.88 |
| 8 | HALLMARK_INFLAMMATORY_RESPONSE | 9.68 | HALLMARK_PANCREAS_BETA_CELLS | 10.78 | HALLMARK_KRAS_SIGNALING_UP | 11.01 | HALLMARK_TNFA_SIGNALING_VIA_NFKB | 8.23 | HALLMARK_TNFA_SIGNALING_VIA_NFKB | 8.20 |
| 9 | HALLMARK_UV_RESPONSE_UP | 9.86 | HALLMARK_E2F_TARGETS | 11.45 | HALLMARK_P53_PATHWAY | 11.39 | HALLMARK_COAGULATION | 8.37 | HALLMARK_EPITHELIAL_MESENCHYMAL_TRANSITION | 8.63 |
| 10 | HALLMARK_E2F_TARGETS | 9.92 | HALLMARK_G2M_CHECKPOINT | 11.51 | HALLMARK_ESTROGEN_RESPONSE_LATE | 11.53 | HALLMARK_KRAS_SIGNALING_UP | 8.40 | HALLMARK_XENOBIOTIC_METABOLISM | 9.22 |
| 11 | HALLMARK_HYPOXIA | 10.64 | HALLMARK_XENOBIOTIC_METABOLISM | 11.68 | HALLMARK_OXIDATIVE_PHOSPHORYLATION | 11.72 | HALLMARK_EPITHELIAL_MESENCHYMAL_TRANSITION | 8.64 | HALLMARK_SPERMATOGENESIS | 9.39 |
| 12 | HALLMARK_G2M_CHECKPOINT | 10.74 | HALLMARK_OXIDATIVE_PHOSPHORYLATION | 11.82 | HALLMARK_HEDGEHOG_SIGNALING | 12.13 | HALLMARK_KRAS_SIGNALING_DN | 8.77 | HALLMARK_KRAS_SIGNALING_UP | 9.56 |
| 13 | HALLMARK_XENOBIOTIC_METABOLISM | 10.78 | HALLMARK_IL6_JAK_STAT3_SIGNALING | 11.89 | HALLMARK_MYC_TARGETS_V1 | 12.17 | HALLMARK_SPERMATOGENESIS | 9.07 | HALLMARK_ESTROGEN_RESPONSE_LATE | 9.59 |
| 14 | HALLMARK_DNA_REPAIR | 10.83 | HALLMARK_MYOGENESIS | 12.40 | HALLMARK_ANDROGEN_RESPONSE | 12.53 | HALLMARK_XENOBIOTIC_METABOLISM | 9.08 | HALLMARK_OXIDATIVE_PHOSPHORYLATION | 9.86 |
| 15 | HALLMARK_ESTROGEN_RESPONSE_LATE | 11.05 | HALLMARK_BILE_ACID_METABOLISM | 12.46 | HALLMARK_APOPTOSIS | 12.53 | HALLMARK_P53_PATHWAY | 9.67 | HALLMARK_P53_PATHWAY | 9.96 |
